# Hfq CLASH uncovers sRNA-target interaction networks involved in adaptation to nutrient availability

**DOI:** 10.1101/481986

**Authors:** Ira A. Iosub, Marta Marchioretto, Brandon Sy, Stuart McKellar, Karen J. Nieken, Rob W. van Nues, Jai J. Tree, Gabriella Viero, Sander Granneman

## Abstract

By shaping gene expression profiles, small RNAs (sRNAs) enable bacteria to very efficiently adapt to constant changes in their environment. To better understand how *Escherichia coli* acclimatizes to changes in nutrient availability, we performed UV cross-linking, ligation and sequencing of hybrids (CLASH) to uncover sRNA-target interactions. Strikingly, we uncovered hundreds of novel Hfq-mediated sRNA-target interactions at specific growth stages, involving many novel 3’UTR-derived sRNAs and a plethora of sRNA-sRNA interactions. We discovered sRNA-target interaction networks that play a role in adaptation to changes in nutrient availability. We characterized a novel 3’UTR-derived sRNA (MdoR), which is part of a regulatory cascade that enhances maltose uptake by (a) inactivating repressive pathways that block the accumulation of maltose transporters and (b) by reducing the flux of general porins to the outer membrane. Our work provides striking examples of how bacteria utilize sRNAs to integrate multiple regulatory pathways to enhance nutrient stress adaptation.

---

Microorganisms are renowned for their ability to adapt to environmental changes by rapidly rewiring their gene expression program. These responses are mediated through integrated transcriptional and post-transcriptional networks. Control at the transcriptional level dictates which genes are expressed (Balleza et al., 2009; Martínez-Antonio et al., 2008) and is well-characterised in *Escherichia coli*. Post-transcriptional regulation is key for controlling adaptive responses. By using riboregulators and RNA-binding proteins (RBPs), cells can efficiently integrate multiple pathways and incorporate additional signals into regulatory circuits. *E. coli* employs many post-transcriptional regulators, including small regulatory RNAs (sRNAs (Waters and Storz, 2009)), *cis*-acting RNAs (Kortmann and Narberhaus, 2012), and RNA binding proteins (RBPs) (Holmqvist and Vogel, 2018). The sRNAs are the largest class of bacterial regulators, which work in tandem with RBPs to regulate their RNA targets (Storz et al., 2011; Waters and Storz, 2009). The base-pairing interactions are often mediated by RNA chaperones such as Hfq and ProQ, which help to anneal or stabilize the sRNA and sRNA-target duplex (Smirnov et al., 2017, 2016; Updegrove et al., 2016). Small RNAs can repress or stimulate translation and transcription, as well as control mRNA stability (Sedlyarova et al., 2016; Updegrove et al., 2016; Vogel and Luisi, 2011; Waters and Storz, 2009).

During growth in rich media, *E. coli* are exposed to continuously changing conditions, such as fluctuations in nutrient availability, pH and osmolarity. Consequently, *E. coli* elicit complex responses that result in physiological and behavioural changes such as envelope composition remodelling, quorum sensing, nutrient scavenging, swarming and biofilm formation. Even subtle changes in the growth conditions can trigger rapid adaptive responses. Accordingly, each stage of the growth curve is characterised by different physiological states driven by activation of different transcriptional and post-transcriptional networks. Moreover, growth phase dependency of virulence and pathogenic behaviour has been demonstrated in both Gram-positive and Gram-negative bacteria. In some cases a particular growth stage is non-permissive for the induction of virulence (Mäder et al., 2016; Mouali et al., 2018). Although the exponential and stationary phases have been characterised in detail(Navarro Llorens et al., 2010; Pletnev et al., 2015), little is known about the transition between these two phases. During this transition, the cell population starts to scavenge alternative carbon sources, which requires rapid remodelling of their transcriptome (Baev et al., 2006a, 2006b; Sezonov et al., 2007).

To understand sRNA-mediated adaptive responses, detailed knowledge of the underlying post-transcriptional circuits is required. In *E. coli*, hundreds of sRNAs have been discovered, but only a fraction of these have been characterised. A key step to unravel the roles of sRNAs in regulating adaptive responses is to identify their targets. To tackle this globally, high-throughput methods have been developed that have uncovered a plethora of sRNA-target interactions, many more than previously anticipated (Han et al., 2016; Hör et al., 2018; Hör and Vogel, 2017; Lalaouna et al., 2015; Melamed et al., 2016; Waters et al., 2016).

To uncover sRNA-target RNA interaction dynamics that take place during the entry into stationary phase, we applied UV cross-linking, ligation and sequencing of hybrids (CLASH) (Helwak et al., 2013; Kudla et al., 2011) on *E. coli*. First, we demonstrate that the highly stringent purification steps make CLASH a robust method for direct mapping of Hfq-mediated sRNA-target interactions in *E. coli*. This enabled us to significantly expand on the sRNA-target interaction repertoire found by RNase E CLASH (Waters et al., 2016) and RIL-seq (Melamed et al., 2016), and we show that Hfq CLASH can generate very reliable results. Using CLASH we identified many potentially novel 3’UTR-derived sRNAs, confirming that this class of sRNAs (Chao et al., 2012, 2017; Chao and Vogel, 2016; Miyakoshi et al., 2015a) is highly prevalent.

Next, we focussed our analyses on interactions that were specifically recovered during the transition phase where we identified a surprisingly large number of interactions, including sRNA-sRNA interactions. Our data suggests that during the transition stage, ArcZ represses CyaR levels, thereby indirectly controlling genes involving nutrient uptake during the transition phase. We also characterized a novel 3’UTR-derived sRNA, which we refer to as MdoR (*mal*-dependent OMP repressor). Unlike the majority of bacterial sRNAs, MdoR is transiently expressed during the transition phase. We demonstrate that MdoR is a degradation intermediate of the *malG* 3’UTR, the last transcript of the *malEFG* polycistron that encodes components of the maltose transport system. We show that MdoR directly downregulates several mRNAs encoding major porins and suppresses the envelope stress response controlled by σ^E^. We propose that MdoR is part of a regulatory network that, during the transition phase, promotes accumulation of high affinity maltose transporters in the outer membrane by repressing competing pathways.

## Results

### Hfq CLASH in *E. coli*

To unravel the post-transcriptional networks that underlie the transition between exponential and stationary growth phases in *E. coli*, we performed CLASH (Helwak et al., 2013; Kudla et al., 2011) using Hfq as bait (Figure 1A). To generate high quality Hfq CLASH data made a number of improvements to the original protocol used for RNase E CLASH (Waters et al., 2016).

**Figure 1.**
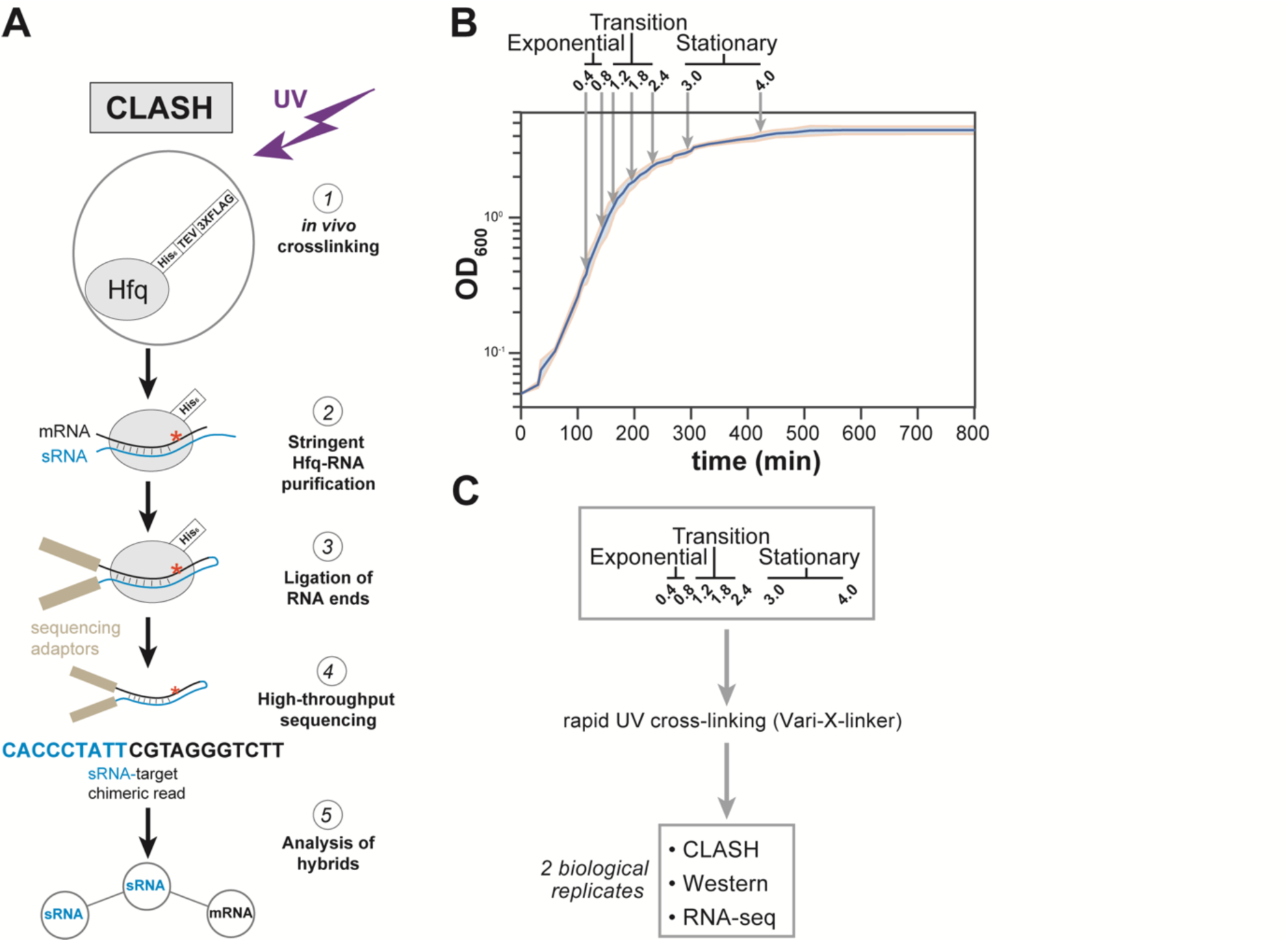
Hfq CLASH experiments at different growth phases in *E. coli*. (**A**) Overview of the critical experimental steps for obtaining the Hfq CLASH data. *E. coli* cells expressing an HTF (His6-TEV-3xFLAG)-tagged Hfq(Jai J. Tree et al., 2014) were grown in LB and an equal number of cells were harvested at different optical densities (OD600). Hfq binds to sRNA-target RNA duplexes, and RNA ends that are in close proximity are ligated together. After removal of the protein, cDNA libraries are prepared and sequenced. The single reads can be used to map Hfq-RNA interactions, whereas the chimeric reads can be traced to sRNA-target interactions. (**B**) A growth curve of the cultures used for the Hfq CLASH experiments, with OD600 at which cells were cross-linked indicated by circles, and each growth stage is indicated above the plot. The results show the mean and standard deviations of two biological replicates. Source data are provided as a Source Data file. (**C**) Cultures at the same OD600 cross-linked and harvested by filtration were analysed by Hfq CLASH, RNA-seq and Western blotting to detect Hfq.

Our Hfq CLASH protocol has several advantages over the related RIL-seq method (see Materials and Methods and Discussion). As negative controls, replicate CLASH experiments were performed on the untagged parental strain. When combined, the control samples had ∼10 times lower number of single-mapping reads and contained only 297 unique chimeric reads, compared to the over 70.000 chimeras identified in the tagged Hfq data. This demonstrates that the purification method produced very low background levels.

Cell samples from seven different optical densities were subjected to Hfq CLASH. Based on the growth curve analysis shown in Figure 1B, we categorized OD_600_ densities 0.4 and 0.8 as exponential growth phase, 1.2, 1.8, 2.4 as the transition phase from exponential to stationary, and 3.0 and 4.0 as early stationary phase. To complement the CLASH data, RNA-seq and Western blot analysis was performed on UV-irradiated cells to quantify steady state RNA and Hfq protein levels, respectively (Figure 1C, Figure 1-figure supplement 1, Supplementary Table 1). Western blot analyses revealed that Hfq levels gradually increased during growth, however, when normalized to the levels of the chaperone GroEL, the increase was modest (Figure 1-figure supplement 1A-B). To determine the cross-linking efficiency, Hfq-RNA complexes immobilized on nickel beads were radiolabelled, resolved on NuPAGE gels and detected by autoradiography. The data show that the recovery of Hfq and radioactive signal was comparable at each optical density studied (Figure 1-figure supplement 1C). Comparison of normalized read counts of replicate CLASH and RNA-seq experiments showed that the results were highly reproducible (Figure 1-figure supplement 2).

### Hfq binds to the transcriptome in a growth-stage dependent manner

Meta-analyses of the Hfq CLASH sequencing data revealed that the distribution of Hfq binding across mRNAs was very similar at each growth stage. We observed the expected Hfq enrichment at the 5’UTRs and at the 3’UTRs at each growth stage (see Figure 1-figure supplement 3A and 3B for examples). After identifying significantly enriched Hfq binding peaks (FDR <= 0.05; see Methods for details) we used the genomic coordinates of these peaks to search for Hfq binding motifs in mRNAs. The most enriched k-mer included poly-U stretches (Figure 1-figure supplement 3C) that resemble the poly-U tracts characteristic to Rho-independent terminators found at the end of many bacterial transcripts (Wilson and Hippel, 1995), and confirms the motif uncovered in CLIP-seq studies in *Salmonella* (Holmqvist et al., 2016).

Given the established role of Hfq in sRNA stabilization and mediating sRNA-target interactions, it was logical to assume that changes in Hfq binding would also be reflected in changes in sRNA steady-state levels. This would imply that the Hfq binding data would show a strong correlation with the RNA-seq data. To test this, we compared the Hfq cross-linking data to the RNA-seq data. K-means clustering of the normalized data revealed 7 different patterns of changes in normalised read counts in the Hfq cross-linking and total RNA-seq data (Figure 2A). Reminiscent of recent work performed in *Salmonella* (Chao et al., 2012), most sRNAs in *E. coli* appear to be preferentially expressed when the cells reach the transition and stationary phase (Figure 2A). However, much to our surprise, the Hfq cross-linking profile did not always follow the same trend (Figure 2A-C, Figure 2-figure supplement 1A). Globally, changes in sRNA expression did not correlate strongly with the Hfq-binding profile (and *vice versa*) (Fig 2B-C). The correlation between changes in sRNA expression levels versus changes in Hfq cross-linking was particularly poor at lower cell density (OD_600_ 0.8: r = 0.10) but gradually improved as the cells approach stationary phase (OD_600_ 4.0 r =0.54; Figure 2B and Figure 2-figure supplement 1A). A very similar result was obtained when comparing all Hfq-bound RNAs, including mRNAs (Figure 2B and Figure 2-figure supplement 1B). Striking examples are MgrR and PsrD, which showed a strong anti-correlation between Hfq CLASH and RNA-seq counts (Figure 2C) In the case of PsrD, Hfq binding showed a modest increase during the growth phase (Figure 2A; right heat map, cluster 6), whereas sRNA levels steadily decreased (Figure 2A; left heat map; cluster 2). PsrD/SraB has also been shown to bind ProQ (Smirnov et al., 2017), which may explain why its accumulation does not correlate with Hfq binding. On the other side of the spectrum, SdsR showed a very high positive Pearson correlation (Figure 2), suggesting that its accumulation heavily relies on Hfq binding.

**Figure 2.**
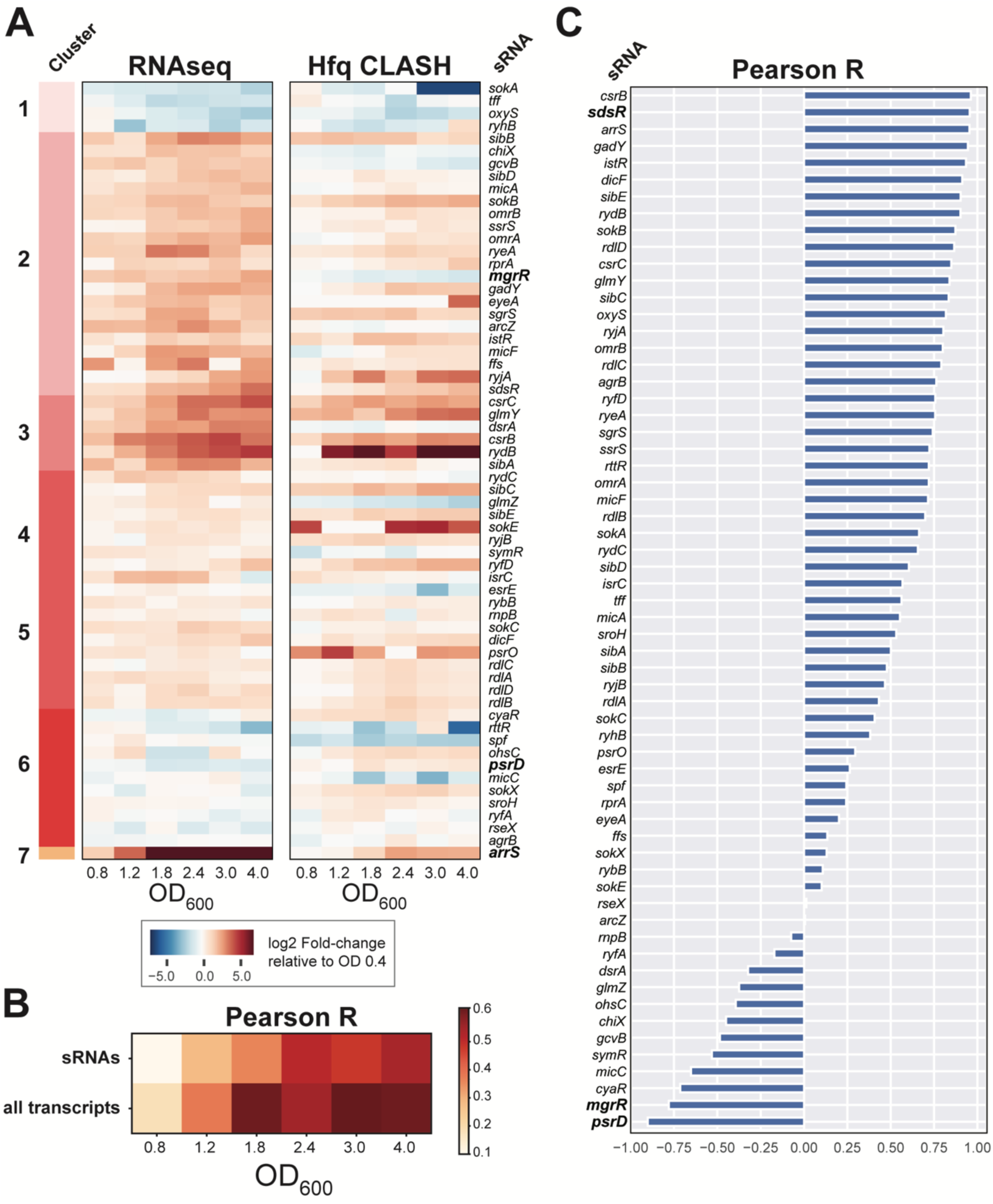
Hfq binding to sRNAs and sRNA steady state levels in *E. coli* do not always correlate highly. (**A**) Heatmaps showing the changes in sRNA steady-states (Left) and Hfq cross-linking (Right) relative to OD600 0.4. The clustered RNA-seq data were generated by k-means clustering using the fold-changes of normalized counts (log2 of transcripts per million (TPM)) relative to OD600 0.4. A blue shade indicates a reduction in levels compared to OD600 0.4, whereas a dark orange shade indicates an increase. The cluster assignment is indicated at the left, and the growth stage is indicated as OD600 units below each heatmap. Note that ssrS and ffs encode the cytoplasmic RNAs 6S, a regulator of RNA polymerase, and 4.5S, the signal recognition particle RNA. EyeA is an uncharacterized sRNA mapped by(Sætrom et al., 2005). (**B**) Global correlation of between Hfq binding and steady state RNA levels increases at higher cell densities. The heatmap shows the changes in Pearson R correlation between Hfq binding and RNA expression (normalised as in (a)), for sRNAs (top) and all transcripts (bottom). (**C**) Assessment of correlation between changes in expression and Hfq binding profiles for individual sRNAs. The y-axis shows the gene name of the sRNAs and the Pearson coefficient indicating the correlation between Hfq binding and steady state levels for each sRNA is shown on the x-axis.

We conclude that the dynamics of sRNA expression and binding to Hfq are not always highly correlated.

### Hfq CLASH robustly detects RNA-RNA interactions

To get a complete overview of the RNA-RNA interactions captured by Hfq CLASH, we merged the data from the two biological replicates of CLASH growth phase experiments (Supplementary Table 2.1). Overlapping paired-end reads were merged and unique chimeric reads were identified using the hyb pipeline (Travis et al., 2013). To select RNA-RNA interactions for further analysis, we applied a probabilistic analysis pipeline previously used for the analysis of RNA-RNA interactions in human cells (Sharma et al., 2016) and adapted it for the analyses of RNase E CLASH data (Waters et al., 2016). This pipeline tests the likelihood that observed interactions could have formed spuriously. Strikingly, 87% of the chimeric reads had a Benjamini-Hochberg adjusted p-value of 0.05 or less, indicating that it is highly unlikely that these interactions were generated by random ligation of RNA molecules. These analyses demonstrate the robustness of Hfq CLASH protocol. A complete overview of statistically significantly enriched chimeras is provided in Supplementary Table 2.2.

The distribution of combinations of transcript classes found in the statistically filtered chimeric reads indicates the sRNA-mRNA interactions as the most frequent recovered Hfq-mediated interaction type (∼25%; Figure 3A). We suspect that this number might be higher, as about 16% of the chimeras contained sRNA and fragments that mapped to intergenic regions (Figure 3A). Manual inspection of several of these indicated that some of the intergenic sequences were located near genes for which the UTRs were unannotated or short. The vast majority of these interactions were growth stage-specific (Figure 3B). Hfq CLASH identified almost 2000 sRNA-mRNA interactions (Figure 3C; Supplementary Table 2). Around 20% of the interactions found with RIL-seq and ∼21% (27 out of 126) of the experimentally verified interactions present in sRNATarbase3 (Supplementary Table 2.7) were recovered. These results suggest that while the CLASH data contained known and many novel interactions, the analyses clearly were not exhaustive.

**Figure 3.**
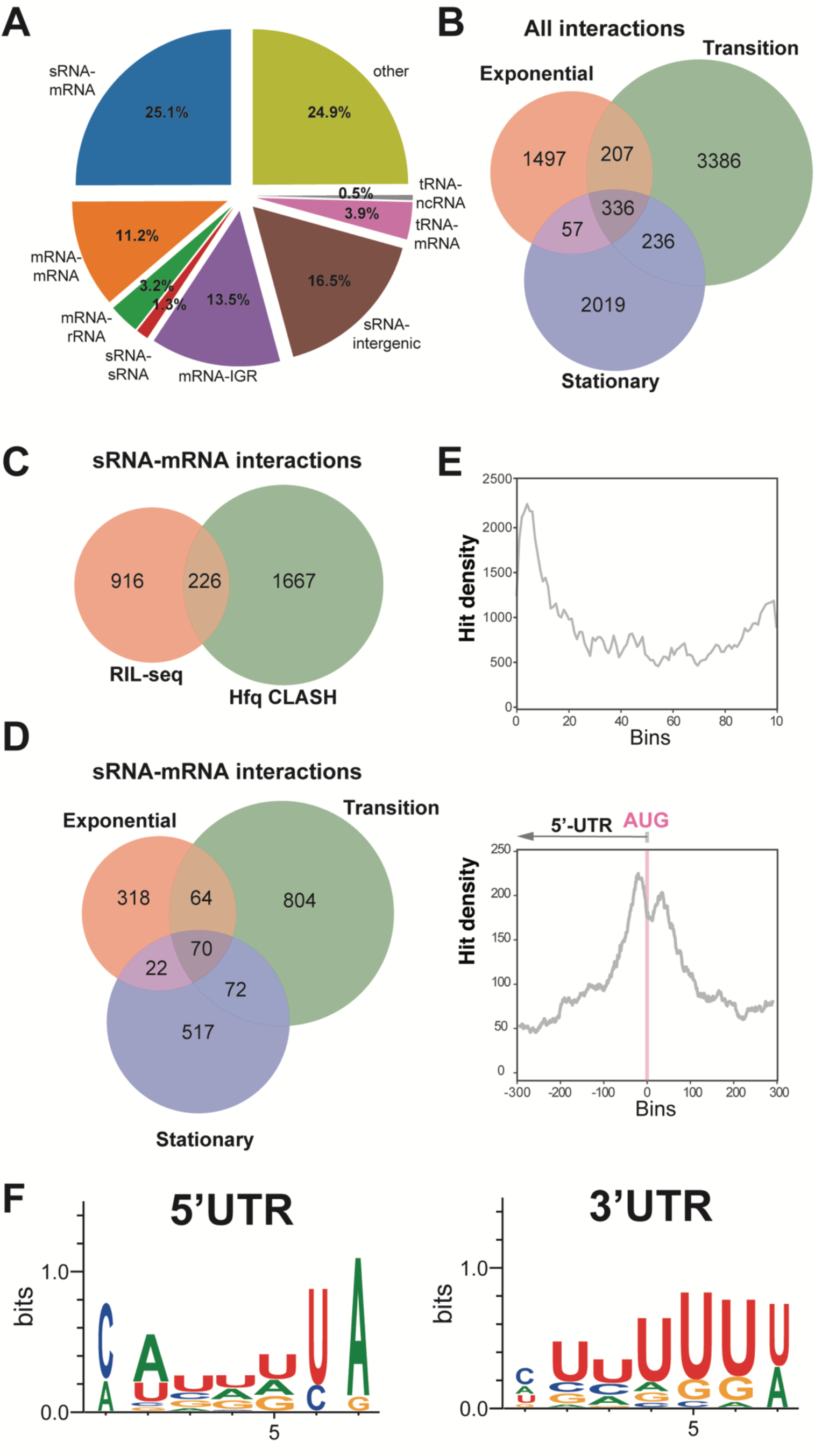
Hfq CLASH detects RNA-RNA interactions in *E. coli*. (**A**) Intermolecular transcript combinations found in interactions captured by Hfq CLASH. Combination count of all uniquely annotated hybrids on genomic features. *tRNA-tRNA and rRNA-rRNA chimeras originating from different coding regions were removed. (**B**) Venn diagram showing the intersection between interactions from statistically filtered CLASH data from two biological replicates, recovered at three main growth stages: exponential (OD600 0.4 and 0.8), transition (OD600 1.2, 1.8, 2.4) and early stationary (OD600 3.0 and 4.0). (**C**) Comparison of sRNA-mRNA interactions found in RIL-seq S-chimera data and Hfq CLASH data. (**D**) Same as in (**B**) but for sRNA-mRNA interactions. (**E**) (Top) Distribution of chimeras representing statistically filtered interactions, which uniquely map to mRNA genes. Overlapping fragments of the two individual parts of the chimeras were collapsed into single clusters followed by generation of distribution plots. Each gene was divided in 100 bins the number of clusters that map to each bin (hit density; y-axis) was calculated; (Bottom) For the distribution plot around the AUG, the gene length was normalized in 601 bins (x-axis) 5’-end overlap (−300) before the start of the coding sequence, and 300 bins downstream AUG (+300); the bins corresponding to the start codon are indicated with a pink line. (**F**) Enriched motifs in chimeras that uniquely overlap 5’UTRs and 3’UTRs; the logos were drawn using the top 20 K-mers.

Meta-analysis revealed that the majority of interactions were identified in the transition phase (Figure 3D) and that the mRNA fragments found in chimeric reads were strongly enriched in 5’UTRs peaking near the translational start codon (Figure 3E). The latter is consistent with the canonical mode of translational inhibition by sRNAs (Bouvier et al., 2008) and demonstrates the robustness of the data. Enrichment was also found in 3’UTRs of mRNAs (Figure 3F). Motif analyses revealed a distinct sequence preference in 5’UTR and 3’UTR binding sites (Figure 3G, Supplementary Table 3). The motifs enriched in the 5’UTRs chimeric fragments are more consistent with Hfq binding to Shine Dalgarno-like (ARN)_n_ sequences (Jai J. Tree et al., 2014) and U-tracts, whereas the 3’UTR-containing chimera consensus motif corresponds to poly-U transcription termination sites (Figure 3F and Supplementary Table 3).

### Hfq CLASH predicts sRNA-sRNA interactions as a widespread layer of post-transcriptional regulation

We also uncovered a surprisingly large number of sRNA-sRNA interactions (Supplementary Table 2.4), many of which were uniquely found in our Hfq CLASH data (Figure 4A). Many interactions were growth-stage specific and the sRNA-sRNA networks show extensive rewiring across the exponential, transition and stationary phases (Figure 4-figure supplement 1). The sRNA-sRNA network is dominated by several abundant sRNAs that appear to act as hubs that have many interacting partners: ChiX, Spot42 (spf), ArcZ and GcvB. In many cases the experimentally-validated sRNA seed sequences were found in the chimeric reads, for both established and novel interactions. For example, the vast majority of ArcZ sRNA-sRNA chimeras contained the known and well conserved seed sequence (Figure 4B, Figure 4-figure supplement 2).

**Figure 4.**
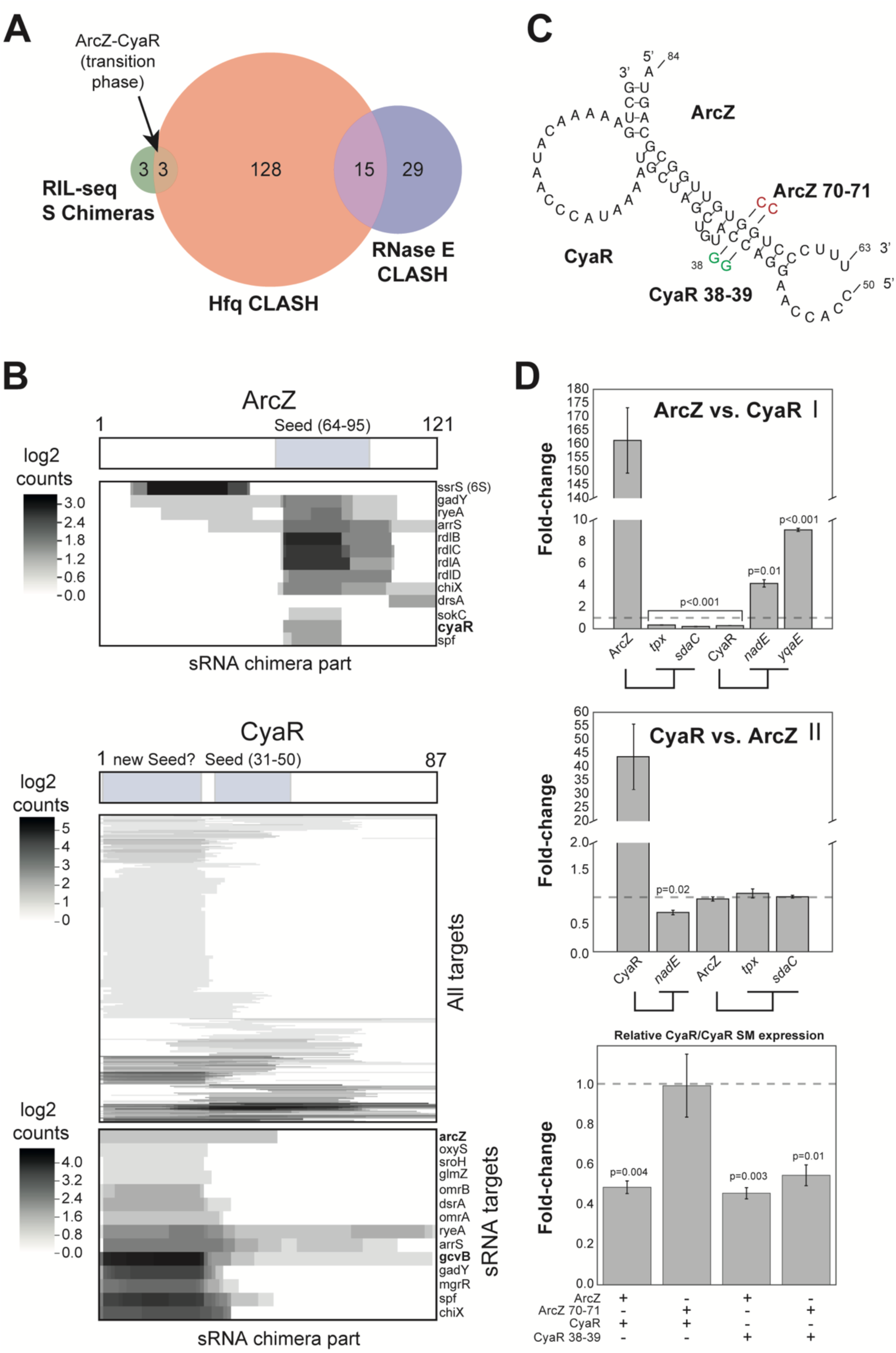
sRNA-RNA interactions identified by CLASH are growth-stage specific. (**A**) Hfq CLASH uncovers sRNA-sRNA networks: comparison between statistically filtered sRNA-sRNA interactions in the Hfq CLASH data, RIL-seq S-chimeras (Melamed et al., 2016) (log and stationary) and RNase E CLASH (Waters et al., 2016). Only core genome sRNAs were considered. Red coloured sRNA-sRNA interactions have been characterized in more detail. (**B**) Heatmaps showing the read density (log2(chimera count)) of chimeric fragments mapping to ArcZ and CyaR. (Top) ArcZ regions involved in sRNA-sRNA interactions. The location of the known ArcZ seed sequence is indicated above. (Bottom) CyaR heatmaps that show all CyaR chimeras and CyaR-sRNA chimeras, respectively. The location of the known CyaR seed sequence, as well as a new seed, is indicated above. (**C**) Base-pairing interactions predicted from the ArcZ-CyaR chimeras using RNAcofold. The nucleotide substitutions for experimental validation of direct base-pairing are shown as red or green residues. (**D**) SRNA-sRNA interactions coordinate nutritional stress responses. ArcZ, CyaR were overexpressed and the levels of their targets were monitored by RT-qPCR. The *tpx* and *sdaC* mRNAs are ArcZ mRNA targets. The *nadE* and *yqaE* mRNAs are CyaR targets. The *dppA* mRNA is a GcvB target. Experiments were performed in biological and technical triplicates; Error bars indicate the standard error of the mean (SEM) of the three biological replicates. The dashed horizontal line indicates the level of the overexpressed scrambled RNA. (**E**) ArcZ and CyaR directly interact. The sRNAs and mutants as in (**C**) were ectopically co-expressed in *E. coli* and CyaR and CyaR 38-39 levels were quantified by RT-qPCR.

The sRNA-sRNA chimeras containing CyaR fragments were of particular interest, as the sRNA is primarily expressed during the transition from late exponential to stationary phase (De Lay and Gottesman, 2009). In the case of CyaR, the known seed sequence (De Lay and Gottesman, 2009; Papenfort et al., 2008) as well as a conserved ∼25 nt fragment in the 5’ region was found in chimeras (Figure 4-C; Figure 4-figure supplement 2). Similar seed sequences were identified in CLASH experiments using RNase E as a bait (Waters et al., 2016), suggesting that this region represents a *bona fide* interaction site. Notably, we identified ArcZ-CyaR chimeras containing the seed sequence from both sRNAs (Figure 4-figure supplement 2) and these were detected specifically in the transition phase (Figure 4B-C), suggesting that these sRNAs could influence each other’s activity. To validate these findings, we used an *E. coli* plasmid-based assay that is routinely used to monitor sRNA-sRNA interactions and expression of their target mRNAs (Melamed et al., 2016; Miyakoshi et al., 2015b; Jai J. Tree et al., 2014). An advantage of this system is that each sRNA would be uncoupled from the chromosomally encoded regulatory networks (that were thought to act largely in a 1:1 stoichiometry) and to allow the specific effects of the sRNA-target RNA to be assessed (Miyakoshi et al., 2015b). Importantly, these sRNAs were induced during early exponential growth phase when the endogenous (processed) ArcZ and CyaR sRNAs are detectable at only very low levels (Figure 4-figure supplement 3B, lanes 1, 2, 5, 7). The qPCR data were subsequently normalized to the results obtained with a control scrambled sRNA to calculate fold changes in expression levels. Since it is difficult to predict directly from the CLASH data which sRNA in each pair acts as the decoy/sponge, we tested both directions. ArcZ overexpression not only decreased the expression of its mRNA targets (*tpx*, *sdaC*) by more than 50% but also that of CyaR (Figure 5D, panel I). Concomitantly, we observed a substantial increase in CyaR targets *nadE* and *yqaE* (Figure 4D, panel I). CyaR overexpression reduced the level of a direct mRNA target (*nadE*) by ∼40% but it did not significantly alter the level of ArcZ or ArcZ mRNA targets (*tpx* and *sdaC*; Figure 4D, panel II). Notably, in this two-plasmid assay CyaR was not expressed at levels higher than ArcZ (Figure 4-figure supplement 3A, panel II). Therefore, it is possible that under the tested conditions the CyaR overexpression was not sufficient to see an effect on ArcZ. We find this unlikely as overexpression of CyaR also did not significantly affect endogenous ArcZ levels, which was ∼80-fold less abundant than CyaR in this experiment (Figure 4-figure supplement 3A, panel III). The qPCR results were also confirmed by Northern blot analyses (Figure 4-figure supplement 3B, lanes 1-8), which also demonstrate that ArcZ processing was not affected upon CyaR overexpression. These results suggest that the regulation is unidirectional, reminiscent of what has been described Qrr3 in *Vibrio harveyi* (Feng et al., 2015). We conclude that ArcZ acts as a CyaR anti-sRNA and can trigger its degradation.

**Figure 5.**
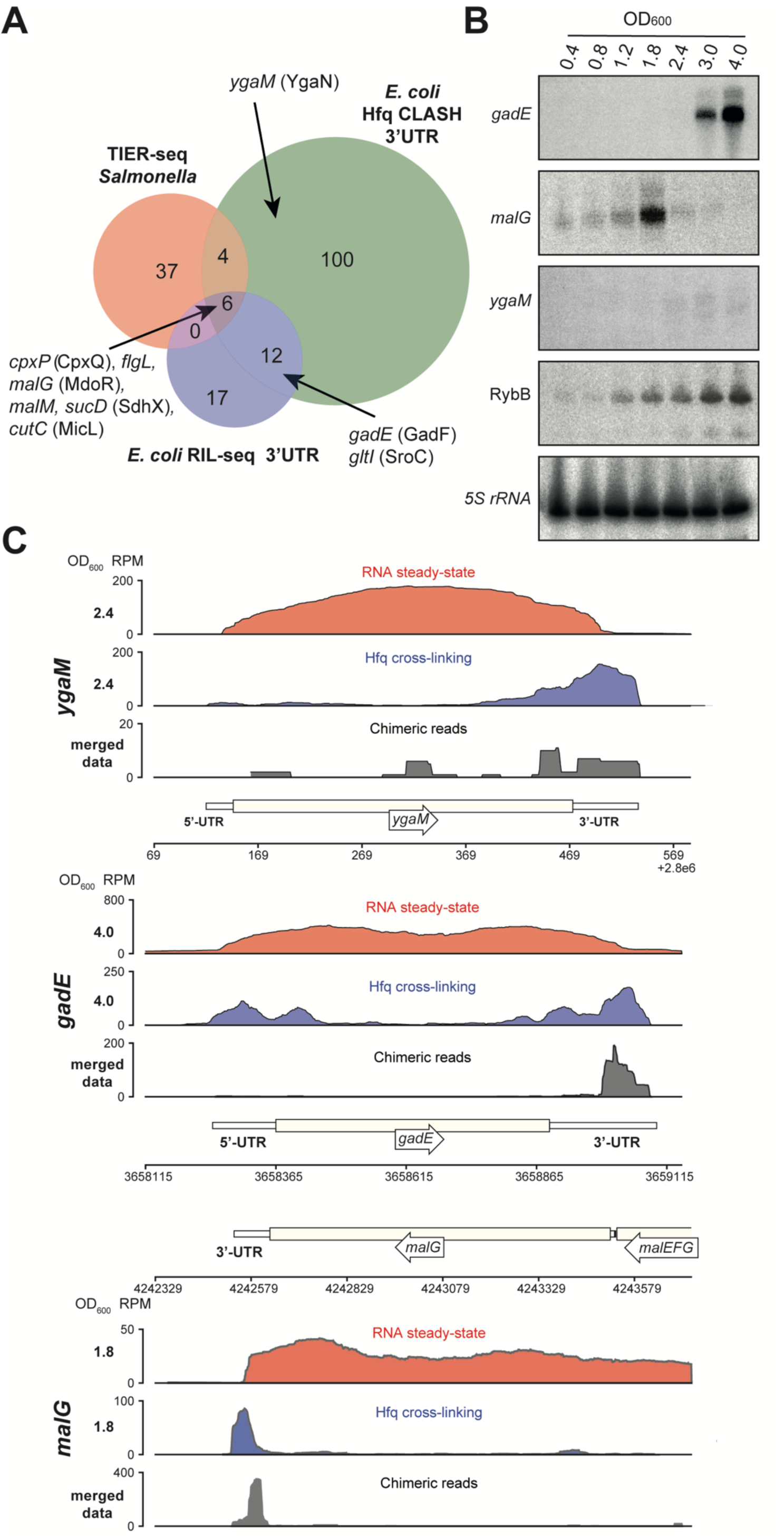
Hfq CLASH uncovers novel 3’UTR-derived sRNAs. (A) Genes of which the 3’UTRs were found fused to mRNAs, were selected from the statistically filtered CLASH data and RIL-seq S-chimera data. The RIL-seq RNA-RNA interaction set (Melamed et al., 2016) S-chimeras for Log and Stationary phases of growth was filtered for the 3’UTR/EST3UTR annotations on either orientation of the mRNA-mRNA pairs. Both were intersected with the set of mRNAs that were predicted by TIER-seq studies (Chao et al., 2017) to harbour sRNAs that get released from 3’UTRs by RNase E processing. Known (CpxQ, SdhX, MicL, GadF and SroC) and novel 3’UTR derived sRNAs (MdoR, *figL* 3’UTR and YgaN) are indicated. (**B**) MdoR is transiently expressed during the transition from exponential to stationary phase. RybB was probed as a sRNA positive control and 5S rRNA as the loading control. See Figure 6-figure supplement 1 for full-size blots. Source data are provided as a Source Data file. (**C**) Genome-browser snapshots of several regions containing candidate sRNAs for optical densities at which the RNA steady-state was maximal for each candidate; the candidate names and OD600 are indicated at the left side of the y-axes; the y-axis shows the normalized reads (RPM: reads per million); red: RPM of RNA steady-states from an RNA-seq experiment, blue: Hfq cross-linking from a CLASH experiment; black: unique chimeric reads found in this region.

To provide additional support for direct interactions between these sRNAs, we generated mutations in the seed sequences of the sRNAs analysed here (Figure 4C). We found that two G to C nucleotide substitutions in ArcZ was sufficient to disrupt ArcZ downregulation of CyaR (Figure 4C-D; ArcZ 70-71 + CyaR). This regulation, however, was almost fully restored when complementary mutations were introduced in the CyaR region (Figure 5C-D; ArcZ 70-71 + CyaR 38-39). These data also demonstrate that it is very unlikely that the observed changes in CyaR levels were be the result of Hfq redistribution due to over-expression of ArcZ over-expression (Moon and Gottesman, 2011; Papenfort et al., 2009), as the ArcZ seed mutant stably accumulated (and therefore effectively binds Hfq), but did not affect CyaR levels. Unexpectedly, the wild-type ArcZ was also able to effectively suppress the CyaR seed mutant (Figure 5D; ArcZ + CyaR 38-39), possibly because ArcZ can still form stable base-pairing interactions with the CyaR mutant.

These results, together with the CLASH data, strongly support the notion that ArcZ and CyaR base-pair *in vivo*, resulting in degradation of CyaR but not *vice versa*.

### Hfq CLASH identifies novel sRNAs in untranslated regions

Two lines of evidence from our data indicate that many more mRNAs may be harbouring sRNAs in their UTRs or be involved in base-pairing among themselves. First, around 11% of the intermolecular chimeras mapped to mRNA-mRNA interactions (Figure 3A). Secondly, we observed extensive binding of Hfq in 3’UTRs near transcriptional terminators (Figure 1-figure supplement 3A, C), indicating that like in *Salmonella*, *E. coli* 3’UTRs may harbour many functional sRNAs (Chao et al., 2017). We identified 122 3’UTR-containing mRNA fragments that were involved in 550 interactions. Sixty-five of these interactions were also identified in the RIL-seq S-chimeras data (Melamed et al., 2016). Eighteen of the 3’UTRs were found as part of chimeras in the RIL-seq data, while 10 appeared stabilised upon transient inactivation of RNase E performed in *Salmonella* (TIER-seq data (Chao et al., 2017)); Figure 6A, Supplementary Tables 2.5 and 2.6). Out of the 550 3’UTR-mRNA chimeric reads, 79 were 3’UTRs fused to 5’UTRs of mRNAs, suggesting that these may represent 3’UTR-derived sRNAs that base-pair with 5’UTRs of mRNAs, a region frequently targeted by sRNAs (Supplementary Table 2.6). Strikingly, 223 interactions contained 3’UTR fragment of *cpxP*, 51 of which were also found in the RIL-Seq data (Supplementary Table 2.6). In *Salmonella cpxP* harbours the CpxQ sRNA (Chao and Vogel, 2016). Our analyses greatly increased the number of potential CpxQ mRNA targets and show that the vast majority of CpxQ interactions take place during the transition and stationary phases (Supplementary Table 2.6).

**Figure 6.**
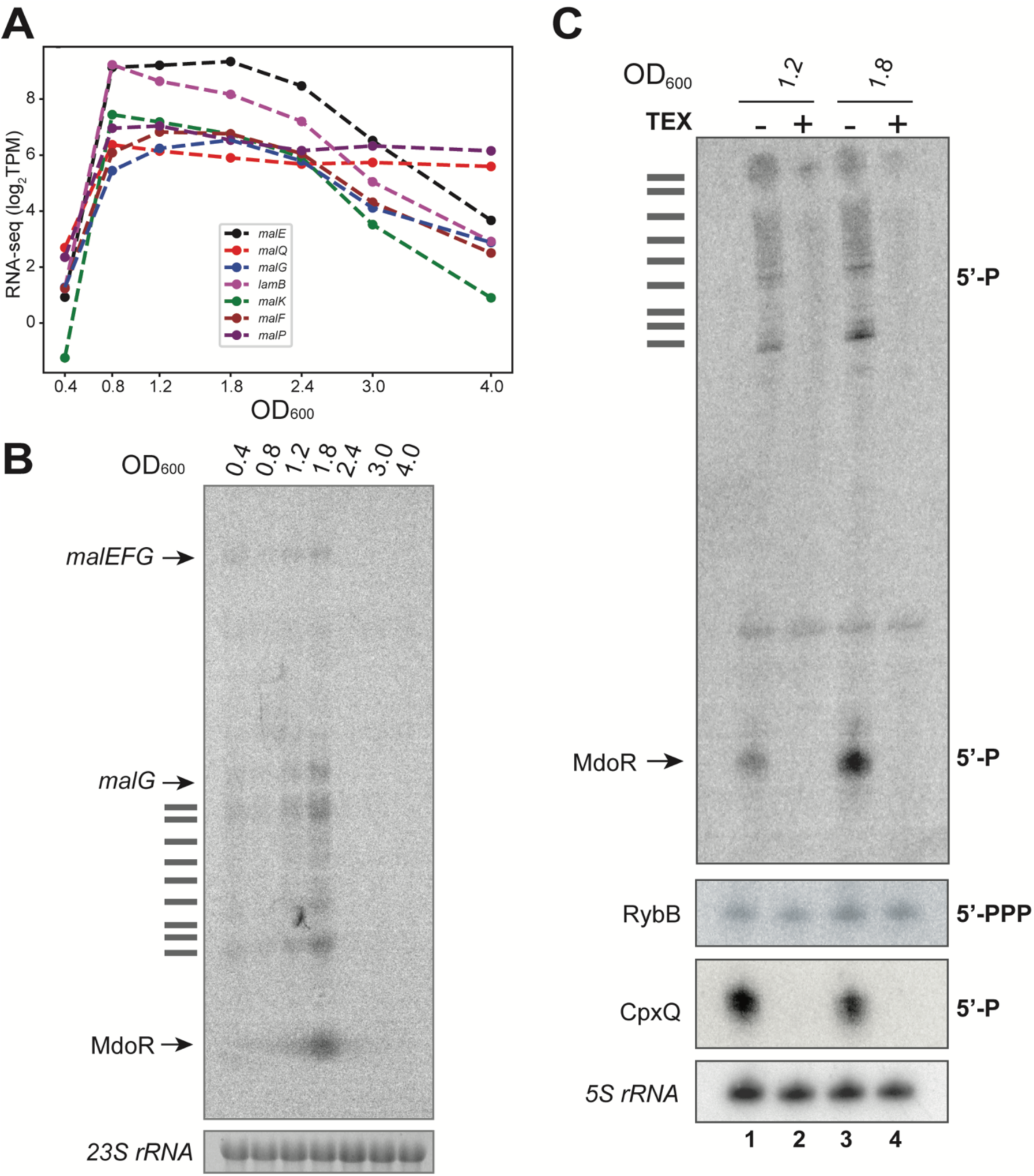
MdoR is a degradation product that emerges at the transition between exponential and stationary phase of growth. (**A**) The *mal* regulon gene expression peaks at the transition between exponential and stationary phases of growth: the plot shows averages of log2(TPM) normalized RNA steady-state levels (y-axis); the x-axis indicates the cell densities (OD600) at which samples were taken. (**B**) Northern blot using total RNA from *E. coli* harvested at different cell densities (OD600) probed with an oligo antisense to *malG* 3’UTR; 23 S rRNA was used as the loading control; the identity of the bands is indicated at the left of the panel; horizontal bars indicate *malEFG* degradation intermediates. (**C**) MdoR is a degradation product: Northern blot using total RNA from cells at indicated OD600 with (+; lanes 2 and 4) or without (-; lanes 1 and 3) 5’-Phosphate-Dependent Exonuclease (TEX) treatment. The sRNAs RybB (5’ppp) and CpxQ (5’p) were used as negative and positive controls, respectively. The 5S rRNA is a loading control. The text on the right of the blot indicates the phosphorylation state of the 5’-termini for each sRNA.

We identified six mRNA 3’UTRs that were uncovered in all three (Hfq CLASH, RIL-seq and TIER-seq) datasets (Figure 5A), suggesting these likely contain sRNAs released from 3’UTRs by RNase E processing. Northern blot analyses confirmed the presence of sRNAs in *malG* and *gadE* 3’UTRs (Figure 5B, Figure 5-figure supplement 1A). The latter was also recently experimentally confirmed in the RIL-seq data and was annotated as GadF (Melamed et al., 2016). Furthermore, significant Hfq cross-linking could be detected in the 3’UTRs of these transcripts (Figure 5C). In addition, we could show that the 3’UTR of *ygaM*, which was found in chimeric reads in our data, also likely harbours a ∼100 nt sRNA (hereafter referred to as YgaN; Figure 5-figure supplement 1A) and robust Hfq cross-linking could be detected in this region (Figure 5B-C).

To substantiate these results, we analysed RNA-seq data from a study that used Terminator 5’-Phosphate Dependent Exonuclease (TEX) to map transcription start sites (TSS) of coding and non-coding RNAs in *E. coli* (Thomason et al., 2015). TEX degrades processed transcripts that have 5’ monophosphates, but not primary transcripts with 5’ triphosphates. Therefore, these data enabled us to determine whether (a) a TSS was detected in the 3’UTR and whether these were generated by RNase-dependent processing (TEX sensitive) or originated from an independent promoter (TEX insensitive). For 47 of the 122 predicted 3’UTR-derived sRNAs TEX data provided strong evidence for the presence of sRNAs (Figure 5-figure supplement 1B-C, Supplementary Table 2.5 and see Data and Code availability). The TEX data indicate that *<u>ygaM</u>* has (at least) two promoters, one of which is located near the 3’ end of the gene that we predict is the TSS for YgaN (Figure 5-figure supplement 1B). Furthermore, we speculate that YgaN is processed by RNases. This is based on the observation that multiple YgaN species were detected in the Northern blot analyses (Figure 5-figure supplement 1A) and the TEX data indicate that shorter YgaN RNAs are sensitive to TEX treatment (Figure 5-figure supplement 1B).

The majority of the sRNAs we analysed are more abundant at higher cell densities (including GadF, YgaN and RybB; see Figure 5B, 2A). In sharp contrast, 3’ UTR *malG* sRNA was expressed very transiently and peaked at an OD_600_ of 1.8 (Figure 5B). We envisage that the particularly transient expression of this sRNA may be correlated with a role in the adaptive responses triggered during transition from exponential to stationary phases of growth. We named it MdoR (*mal*-dependent OMP repressor) and characterized it in detail.

The steady state levels of *malG* and *malEFG* transcripts recapitulate the same expression profile: both peak at OD_600_ of 1.8 and drop to very low levels at OD_600_ 2.4 (Figure 6A). Additionally, we identified shorter *malG* 3’UTR-containing fragments of intermediate length between *malG* and MdoR that could be degradation intermediates (Figure 6B). The detection of these intermediate species suggests that the *malEFG* primary transcript is undergoing serial ribonucleolytic cleavage steps.

MdoR is a 104 nt sRNA that contains part of the *malG* coding sequence, including the stop codon and the Rho-independent terminator (Figure 5-figure supplement 1A-B). Two lines of evidence suggest that MdoR is generated via endonucleolytic cleavage: RNase E cleavage was detected in the 3’UTR of *malG* in the *Salmonella* TIER-seq data (Chao et al., 2017). Secondly, the TEX RNA-seq data supported the existence of a short RNA in the same region that has a 5’ monophosphate (Figure 5-figure supplement 1B-C). We verified the MdoR TEX data by Northern blot analyses (Figure 6C). Consistent with the TEX RNA-seq data, MdoR could not be detected in our RNA samples treated with TEX (Figure 6C, lanes 2 and 4), confirming it bears a 5’ monophosphate. Like MdoR, the positive control CpxQ (Chao and Vogel, 2016) was degraded in the presence of TEX (Figure 6C). In contrast, RybB, an sRNA with a 5’ triphosphate transcribed from an independent promoter (Johansen et al., 2006; Papenfort et al., 2006) was a poor substrate for the exonuclease (Figure 6C).

TEX treatment of the total RNA also reduced the levels of the longer intermediate species as well as the full-length *malG*. These data support a mechanism by which the full-length polycistronic RNA undergoes decay that is initiated at a site in the upstream *malEFG* region. The distal gene *malE* is clipped off by the degradosome and selectively stabilized, allowing it to be expressed at higher levels than other members of the operon (Newbury et al., 1987). The *malG* 3’UTR, however, would be less susceptible to degradation as it is stabilized by Hfq binding.

### MdoR directly regulates the expression of major outer membrane porins and represses the envelope stress response pathway

The MdoR chimeras frequently contained 5’UTR fragments of two mRNAs encoding major porins, *ompC* and *ompA* (Figure 7A-C), which were also significantly enriched in the RIL-seq data (Figure 8B). The Hfq CLASH data, however, also contained MdoR fragments fused to a *cis*-encoded sRNA, OhsC (Figure 7B-C), suggesting that either MdoR controls its expression by sponging/degradation, or *vice versa*). The most abundant and favourable interactions of MdoR with mRNAs (*ompC* and *ompA*) appear to be utilizing roughly the same region of the *malG* 3’UTR for base-pairing (Figure 7C), suggesting that the corresponding site on the predicted sRNA may be a main, functional seed. The predicted interaction between MdoR and *ompC* is unusually long and consists of two stems interrupted by a bulge, suggesting that these two RNAs form a stable complex. Conservation analyses and *in silico* target predictions (CopraRNA (Wright et al., 2014, 2013)) indicate that the seed sequence predicted by CLASH is relatively well-conserved (Figure 7-figure supplement 1A), and could be utilized for the regulation of multiple targets (Figure 7-figure supplement 1B).

**Figure 7.**
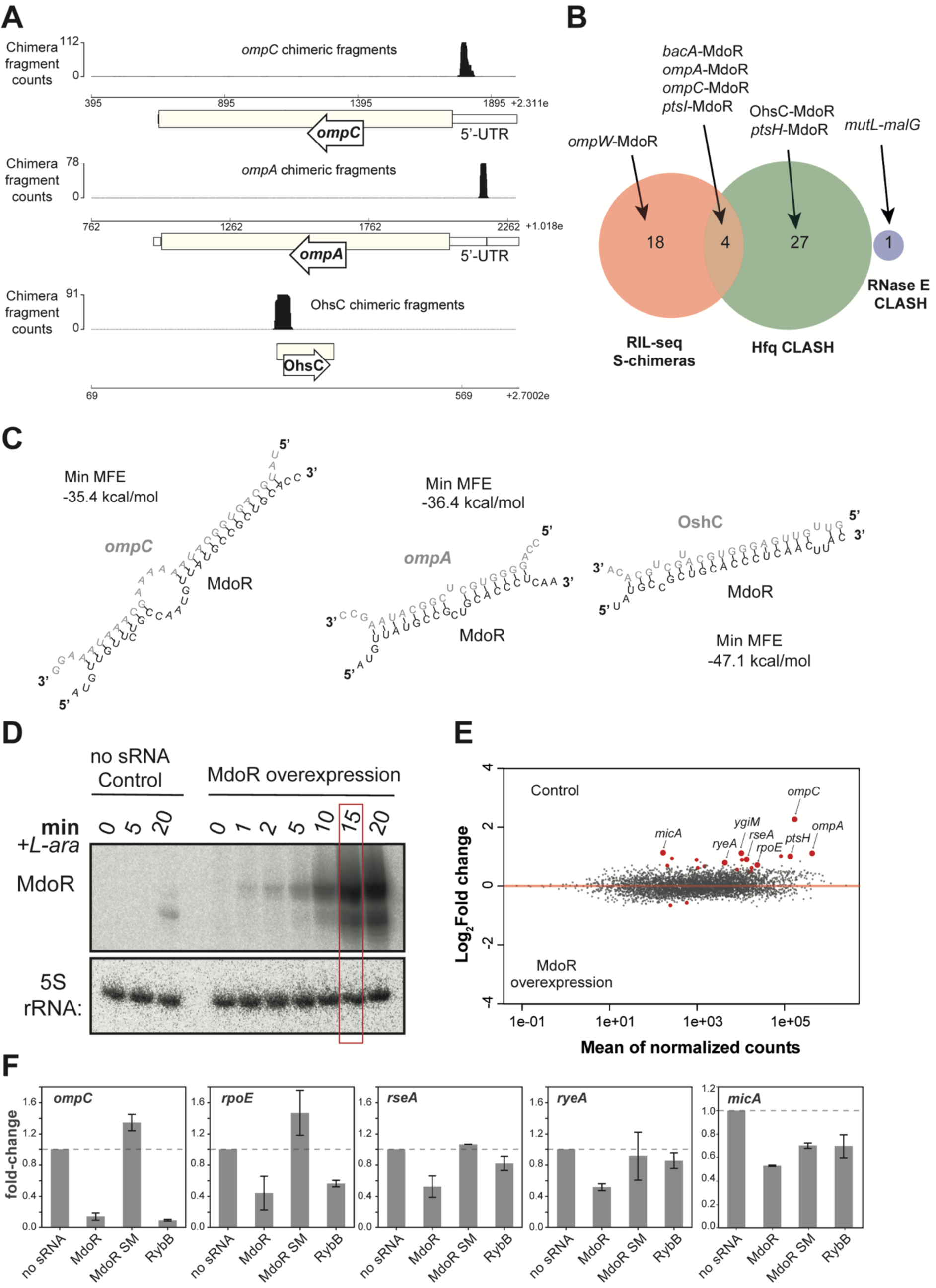
MdoR destabilizes *ompC* and *ompA* mRNAs and downregulates key members the σ^E^ regulon. (**A**) The three genome browser tracks show the distribution of target mRNA and sRNA fragments (*ompC, ompA* and OhsC) that were fused to MdoR fragments in chimeric reads. (**B**) MdoR-target interactions found in Hfq CLASH, RIL-seq S-chimera data (log and stationary phase) and RNase E CLASH data. (**C**) MdoR forms stable duplexes with the 5’UTR of porin-encoding mRNAs and the OhsC sRNA. *In silico* prediction (RNAcofold (Lorenz et al., 2011)) of hybrid structures derived from the most abundant MdoR chimeric reads with the indicated transcripts. The min. MFE is the minimum folding energy assigned by RNAcofold. (**D**) Pulse-overexpression of MdoR using L-arabinose. The empty pBAD plasmid served as a negative control. Samples were harvested 15-minutes after induction. (**E**) MdoR regulates expression of various mRNAs. DESeq2 analyses were performed on RNA-seq data from three biological replicates. Red points indicate differentially expressed transcripts. Transcripts with a log2 fold-change > 0 were enriched in the Control data; those with a log2 fold-change < 0 were enriched in the MdoR overexpression data. The annotated, enlarged red data points indicate several differentially expressed transcripts discussed in the text. (**F**) The MdoR seed region is important for target regulation. RT-qPCR analysis of several differentially expressed transcripts (gene names shown at the top of the plots), in the presence of sRNAs. The plasmid-borne sRNAs were induced for 15 minutes using L-arabinose. The sRNA names are indicated below each bar; the ‘no sRNA’ sample contains the empty plasmid as reference for fold-change calculations; *recA* was used as the internal reference gene; experiments were performed in technical triplicates; the standard error of the mean (SEM) of two biological replicates are reported as error bars.

**Figure 8.**
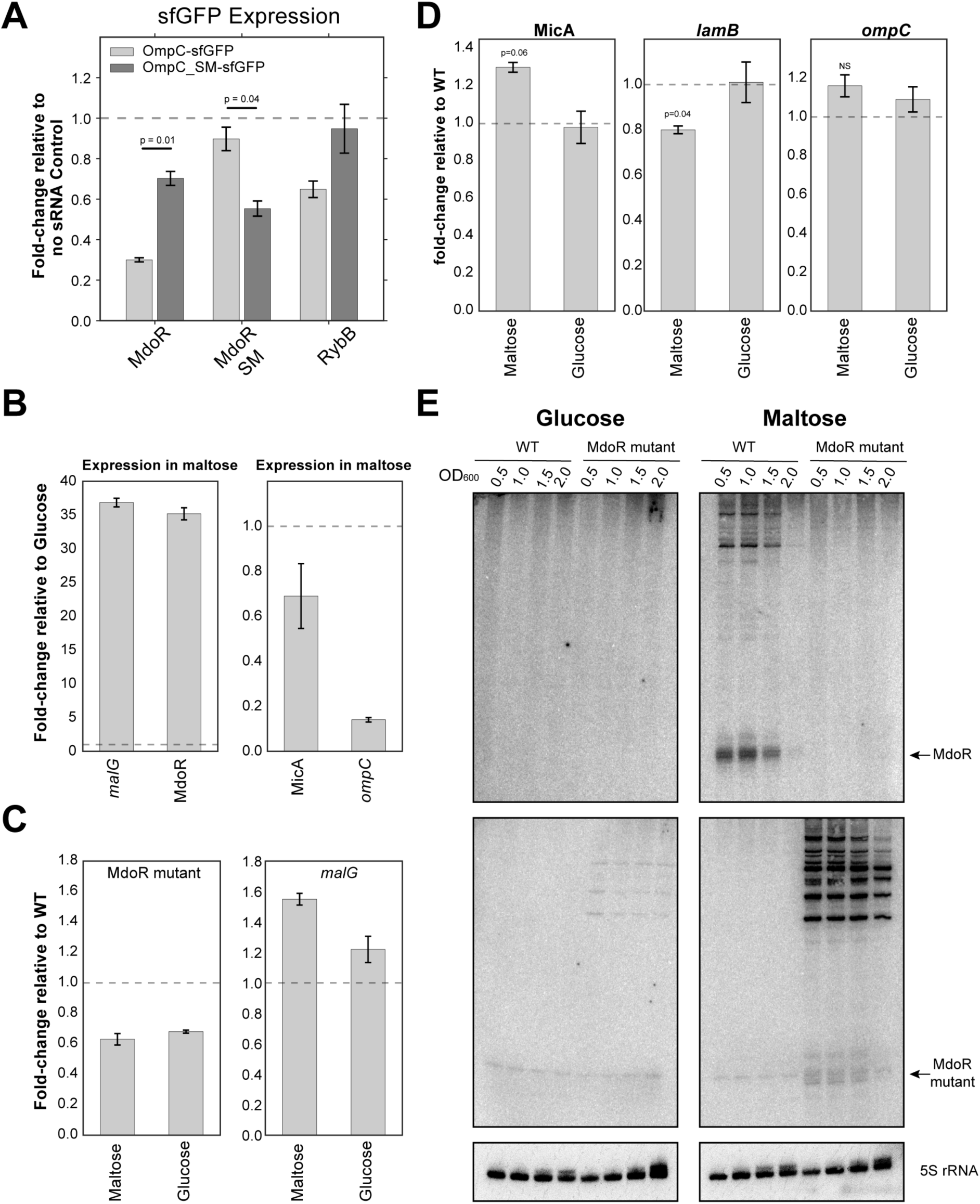
MdoR modulates expression of key factors involved in maltose intake. (**A**) MdoR directly downregulates *ompC* mRNA through base-pairing interactions: RT-qPCR analyses of the *ompC-sfGFP* and *ompC SM-sfGFP* fusions expression in the presence of MdoR. The bars indicate the mean fold-change in expression relative to the no sRNA Control (horizontal dashed line). Error bars indicate the standard error of the mean from two biological replicates. The significance of the differences between the WT and MdoR SM was assessed with a two-tailed Student’s t-test. (**B**) Endogenous MdoR and *malG* expression is induced during growth on maltose, and *ompC* levels are significantly lower in maltose compared to glucose. Total RNA extracted from exponentially growing cells (OD600 0.5) and MdoR, *malG* (Left), and MdoR direct (*ompC*) and indirect (MicA) targets was quantified by RT-qPCR. The data were normalized to 5S rRNA levels. The bars indicate the mean fold-change in expression relative to expression in cells growing in glucose (indicated on the plot with a horizontal dashed line). Error bars indicate the standard error of the mean from two biological replicates. (**C**) The mutant MdoR is ∼50% less abundant than the wild-type. Cell growth and RT-qPCR analysis of MdoR and *malG* expression was performed as in (b). The bars indicate the mean fold-change in expression relative to the wild-type. Error bars indicate the standard error of the mean from two biological replicates. (**D**) Increased MicA and decreased *lamB* levels in the MdoR seed mutant. P-values were calculated using with a one-sample t-test. (**E**) MdoR seed sequences are important for RNase E recruitment and MdoR biogenesis. Northern blot that compares MdoR and longer *malG*-3’UTR containing fragments expression in wild-type *E. coli* and the MdoR seed mutant strain.

To verify the MdoR CLASH data we pulse-overexpressed the sRNA from a plasmid-borne arabinose inducible promoter followed by RNA sequencing (Figure 7D-E). To minimize secondary changes in gene expression, cells were harvested after only 15 minutes of MdoR induction. Note that induction was performed at OD_600_ = 0.4, when endogenous levels of MdoR are very low. As a control we used cells harbouring an empty vector. Differential gene expression analysis (DESeq2 (Love et al., 2014)), identified ∼20 transcripts that were significantly enriched in the control data compared to the MdoR overexpression data (Figure 7E-F; Supplementary Table 4). Thus, these transcripts are likely downregulated by MdoR *in vivo*. This set of transcripts included the sigma factor *rpoE* (σ^E^), which plays an important role in controlling gene expression during stress responses, including envelope stress (Alba and Gross, 2004; Bossi et al., 2008; De Las Peñas et al., 1997; Rhodius et al., 2006). The observed reduction in mRNA levels of the anti-σ^E^ protein RseA can be explained by the fact that σ^E^ and RseA are encoded by the same operon.

Intriguingly, MdoR overexpression also reduced the levels of sRNAs RyeA and MicA, the latter of which depends on σ^E^ for its expression (Udekwu and Wagner, 2007). In *Salmonella*, MicA downregulates LamB, a high affinity maltose/maltodextrin transporter (Bossi and Figueroa-Bossi, 2007). Fragments of three mRNAs (*ompC*, *ompA* and *ptsH*) that were found in MdoR chimeric reads were also differentially expressed in the MdoR overexpression RNA-seq data, providing strong evidence that these are direct MdoR targets.

All the available data suggest that MdoR is part of a mixed coherent feed forward regulatory network (FFL) that enhances the uptake of maltose/maltodextrin by maltose transporters (see Discussion). By base-pairing with the 5’UTRs of the mRNAs it firstly reduces the flux of the more general porins such as OmpC and OmpA to the outer membrane. Secondly, we propose that MdoR stimulates the accumulation of the high-affinity maltose porin LamB in the OMP by suppressing the inhibitory σ^E^ pathway and MicA. To further test this model, we performed additional validation experiments. First, we confirmed the DESeq2 results for a number of the regulated genes (*ompC, rpoE*, *micA* and *ryeA*) by RT-qPCR (Figure 7F). We included an MdoR mutant in which seed sequence in stem 1 was changed into its complementary sequence (Figure 8-figure supplement 1A). As control we also included the RybB sRNA, which regulates *rpoE* and *ompC* expression (Gogol et al., 2011; Papenfort et al., 2006; Thompson et al., 2007). In all cases, the MdoR SM mutations reduced the negative regulatory effect on target expression (Figure 7F).

To demonstrate direct target regulation *in vivo*, we employed a well-established reporter system where an sRNA is co-expressed with a construct containing the mRNA target region fused to the coding sequence of superfolder green fluorescent protein (sfGFP) (Corcoran et al., 2012; Urban and Vogel, 2007) (Figure 8-figure supplement 1B). Fusions for *ompC*, *ompA* and σ^E^ were constructed, but only the OmpC and OmpA-sfGFP reporters produced stable fusions that could be analysed. We also included an MdoR sRNA seed sequence mutant (MdoR SM) and an *ompC* mutant containing compensatory mutations (OmpC SM; Figure 8-figure supplement 1A). Unfortunately, our OmpA-GFP reporter construct that contained compensatory mutations in the target region did not generate a stable fusion. Therefore, we were unable to use this reporter system to verify the MdoR-*ompA* interaction. As positive controls we used the MicC and RybB sRNAs as they both regulate *E. coli ompC* expression (Chen et al., 2004; Gogol et al., 2011). Fluorescence measurements confirmed that levels of OmpC-sfGFP and OmpA-sfGFP fusions were significantly lower in cells expressing MdoR (Figure 8-figure supplement 1C). Importantly, MdoR overexpression did not change the expression of the GFP reporter itself (Figure 8-figure supplement 1C). Mutating the MdoR seed region largely restored OmpA- and OmpC-sfGFP reporter levels to the levels of the no sRNA negative control. The MdoR SM mutant was still able to partially suppress the expression of the OmpC SM-sfGFP mutant, suggesting that base-pairing interactions between these two mutants is less stable compared to the wild-type (Figure 8-figure supplement 1C). The wild-type MdoR was also able to partially suppress the expression of the *ompC* SM mutant. We suggest that the predicted base-pairing interactions between MdoR and *ompC* in the second stem (Figure 8-figure supplement 1A) might be sufficient to partially suppress *ompC* expression. Regardless, the data strongly imply that MdoR directly regulates *ompC* expression.

To determine whether the changes in fluorescence signal correlate with changes in reporter mRNA levels, we measured the expression of the GFP reporters by RT-qPCR. The results were essentially identical to the GFP fluorescence measurements (Figure 8A); Overexpression of the wild-type MdoR, but not seed mutant, reduced *ompC*-sfGFP mRNA levels. The *ompC* seed mutation (SM) did not fully disrupt regulation by wild-type MdoR. However, the MdoR mutant containing compensatory mutations (SM mutant) was able to much better suppress the *ompC* SM mRNA levels, consistent with the idea that base-pairing was (largely) restored. Next, we performed polysome profiling experiments to assess the level of *ompC* translation upon overexpression of MdoR. Although MdoR overexpression did not noticeably affect 70S and polysome levels (Figure 8-figure supplement 2A), we observed a significant (∼37%) reduction of *ompC* mRNA in the polysomal fractions, relative to the upper fractions (Figure 8-figure supplement 2B). We conclude that MdoR can regulate *ompC* expression at the post-transcriptional and translational level.

### MdoR enhances maltoporin expression during maltose fermentation

To further substantiate these results, we next switched to a more controlled system to investigate the effect of endogenous MdoR on its targets. To determine whether MdoR has a role in adaptation to maltose-metabolising conditions, single overnight cultures grown in glucose were split and (re)inoculated in fresh medium containing either glucose or maltose as the sole carbon sources, and expression of several *mal* regulon genes and MdoR targets were quantified. We show that MdoR and its parental transcript *malG* are almost undetectable during growth in glucose, and highly expressed during growth in maltose (∼35-fold increase, Figure 8B). This is consistent with catabolite repression of the *mal* regulon by glucose, and its induction my maltose (Boos and Shuman, 1998). Intriguingly, we observed that *ompC* mRNA levels are overall significantly lower during growth in maltose, compared to glucose (Figure 8B). This suggests that porin expression is also regulated by nutrient source in *E. coli*. Similarly, MicA, a repressor of LamB synthesis, has reduced levels in maltose compared to glucose (Figure 8B). We next mutated the entire seed sequence of the chromosomal copy of MdoR (both stem 1 and 2; Figure 8-figure supplement 1A) in the chromosomal context, to completely disrupt base-pairing with *ompC* mRNA. Notably, the fully-processed mutant MdoR sRNA is less abundant than the wild-type (Figure 9C) and longer (unprocessed) fragments that contain upstream *malG* regions could be readily detected (Figure 9E). We speculate that the mutation in the upstream MdoR sequence might have affected RNase E recruitment or cleavage, impairing MdoR processing. The MdoR mutant strain also accumulated significantly higher levels of MicA (Figure 8D) and less *lamB*. Levels of *ompC* mRNA levels were also slightly elevated in the mutant in maltose relative to the parental strain, although in this case a similar increase was also observed in glucose.

**Figure 9.**
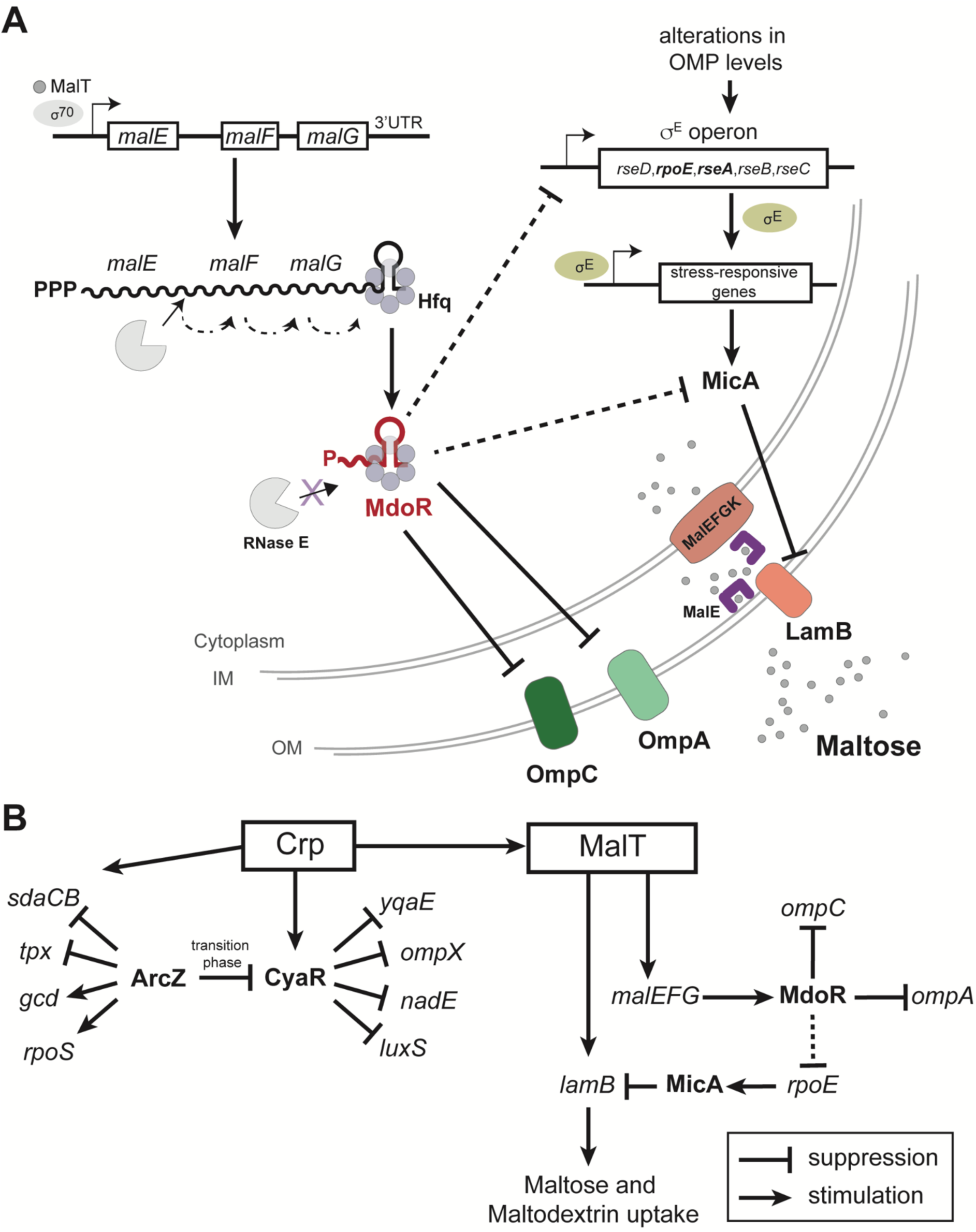
CLASH uncovers sRNA-target interaction networks that regulate adaptation to changes in nutrient availability. (**A**) Model for MdoR biogenesis and its role in post-transcriptional regulation. During maltose (grey circles) utilization, MalT transcribes the *malEFG* operon that encodes components for maltose uptake. A fraction of the *malEFG* transcripts is degraded by RNase E, generating the 3’UTR-derived MdoR sRNA. MdoR base-pairs with *ompC* mRNA, leading to its decay. Additionally, MdoR downregulates expression of *ompA* and MicA that represses LamB synthesis. Therefore, as part of a feed-forward loop (FFL), MdoR promotes accumulation of maltose specific porins, which facilitates maltodextrin uptake. Increased production of LamB exerts additional pressure on the OM assembly machinery. This extracytoplasmic stress, is suppressed by σ^E^ activation, that controls the expression of stress-responsive genes and most of the assembly and insertion factors. But σ^E^ activation is not desirable in these conditions for two reasons: its unbalanced activation has precarious effects on the cell, and would produce high amounts of MicA, that would ultimately repress LamB production. For these reasons, MdoR regulation includes a regulatory arm that mitigates excessive σ^E^ response through MdoR repression of *rpoE* expression and prevention of σ^E^ activation. The latter is achieved by downregulation of *omp* mRNAs, which indirectly relieves envelope stress. (**B**) sRNA-target networks regulating peptide/amino acids and maltose/maltodextrin uptake. Boxed are the key transcription factors, sRNAs are in bold and italicized names are mRNA targets. CRP-cAMP induces expression of CyaR, which is repressed by ArcZ, specifically during the transition phase, a regulatory circuit that connects multiple pathways related to the onset of stationary phase and biofilm formation to quorum sensing, cellular adherence and the nutritional state of the cell. MalT transcribes *lamB* and, via *malEFG* transcription, promotes MdoR accumulation. MdoR indirectly promotes LamB synthesis by repressing the opposing σ^E^ pathway and its sRNA, MicA. Thus, MalT and MdoR jointly promote *lamB* expression, forming a mixed coherent FFL.

Collectively, the data suggest a role for MdoR in enhancing the uptake of maltose when more favourable carbon sources become limiting.

## Discussion

Microorganisms need to constantly adapt their transcriptional program to counteract changes in their environment, such as changes in temperature, cell density and nutrient availability. In bacteria, small RNAs (sRNAs) and their associated RNA-binding proteins are believed to play a key role in this process. By controlling translation and degradation rates of mRNAs upon stress imposition (Holmqvist and Wagner, 2017; Nitzan et al., 2017; Shimoni et al., 2007), they can regulate the kinetics of gene expression as well as suppress noisy signals (Beisel and Storz, 2011), enabling organisms to more efficiently adapt to environmental changes. A major challenge for bacteria is the transition from exponential growth to stationary phase, when the most favourable nutrients become limiting. To counteract this challenge, cells need to rapidly remodel their transcriptome to be able to efficiently metabolize alternative carbon sources. This transition is very dynamic and involves activation and repression of diverse metabolic pathways. However, it is unclear to what degree sRNAs contribute to this transition. The most useful piece of information would be to know what sRNAs are upregulated during this transition phase and to identify their RNA targets. This would help to uncover the regulatory networks that govern this adaptation, as well as provide a starting point for more detailed functional analyses on sRNAs predicted to play a key role in this process. For this purpose, we performed UV cross-linking, ligation and sequencing of hybrids (CLASH (Kudla et al., 2011)) to unravel the sRNA-target interactions during this transition. Using Hfq as a bait we uncovered thousands of unique sRNA-target interactions. Our data are consistent with previously published work (Melamed et al., 2016; Waters et al., 2016) but we also identified almost 1700 novel sRNA-mRNA interactions and over 100 novel sRNA-sRNA interactions. We experimentally validated several of the interactions found in our CLASH findings. We identified functional sRNA-sRNA interactions and describe a novel 3’UTR derived sRNA that plays a role in enhancing uptake of an alternative carbon source during the transition to stationary phase.

### Hfq CLASH

Our *S. cerevisiae* Cross-linking and cDNA analysis data (CRAC; (Granneman et al., 2009)) showed that a percentage of the cDNAs were formed by intermolecular ligations of two RNA fragments (chimeras) known to base pair *in vivo* (Kudla et al., 2011). These findings prompted us to develop a refined protocol to enrich for sRNA-target chimeric reads using Hfq as an obvious bait. The initial Hfq UV cross-linking data (CRAC; (Tree et al., 2014)) did not yield sufficiently high numbers of chimeric reads to extract new biological insights. In line with observations from other groups (Bandyra et al., 2012; Bruce et al., 2018; Morita et al., 2005), it was proposed that duplexes formed by Hfq are rapidly transferred to the RNA degradosome (Bandyra et al., 2012; Bruce et al., 2018; Morita et al., 2005). This can cause an extensive reduction in the likelihood of capturing sRNA-target interactions with Hfq using CLASH (Waters et al., 2016). However, a recent study demonstrated that Hfq can be used effectively as a bait to enrich for sRNA-target duplexes under lower-stringency purification conditions suggesting that sRNA-mRNA duplexes are sufficiently stable on Hfq during purification (Melamed et al., 2016). This encouraged us to further optimize the CLASH method. We made a number changes to the protocol that, when combined, enabled us to recover a large number of sRNA-target chimeric reads (detailed in Materials and Methods). We shortened various incubation steps to minimize RNA degradation and performed very long and stringent washes after bead incubation steps to remove background binding of non-specific proteins and RNAs. Crucially, we very carefully controlled the RNase digestion step that is used to trim the cross-linked RNAs prior to making cDNA libraries, ensuring the recovery of longer chimeric RNA fragments. The resulting cDNA libraries were paired-end sequenced to increase the recovery of chimeric reads with high mapping scores from the raw sequencing data. These modifications led to a substantial improvement in the recovery of chimeric reads (9.5% compared to 0.001%. 0.71% were intermolecular chimeras).

Both RIL-seq and Hfq CLASH have advantages and disadvantages, but they are highly complementary approaches. A major strength of CLASH, however, is that the purification steps are performed under highly stringent and denaturing conditions. During the first FLAG affinity purification steps the beads are extensively washed with high salt buffers and the second Nickel affinity purification step is done under completely denaturing conditions (6M guanidium hydrochloride). These stringent purification steps enable can significantly reduce noise by strongly enriching for RNAs covalently cross-linked to the bait protein (Granneman et al., 2009). Indeed, we show that Hfq CLASH can generate high quality RNA-RNA interaction data with very low background: Only a few hundred chimeric reads were found in control datasets, compared to the over 70.000 that co-purified with Hfq. The RIL-seq library preparation protocol uses an rRNA depletion step to remove contaminating ribosomal RNA, whereas for Hfq CLASH this is not necessary. Our library preparation protocol also includes the use of random nucleotides in adapter sequences to remove potential PCR duplicates (“collapsing”) from the data.

The very stringent purification conditions used in CLASH could, in some cases also be a disadvantage as it completely relies on UV cross-linking to isolate directly bound RNAs. In cases where protein-RNA cross-linking efficiencies are low (for example proteins that only recognize double-stranded RNA), RIL-seq may be a better approach as it does not completely rely on UV cross-linking (Melamed et al., 2016).

A large number of interactions were unique to both RIL-seq and Hfq CLASH datasets, which we believe can be explained by a number of technical and experimental factors. The denaturing purification conditions used with CLASH completely disrupts the Hfq hexamer ((Tree et al., 2014) and this work) and therefore during the adapter ligation reactions the RNA ends are likely more accessible for ligation. In support of this, in the RIL-Seq data, the sRNAs are mostly found in the second half of the chimeras (Melamed et al., 2016), in the Hfq CLASH data we see sRNAs fragments roughly equally distributed in both sides (45% in left fragment and 55% in right fragment). Indeed, it was proposed that in RIL-seq the 3’ end of the sRNA is buried in the hexamer and therefore not always accessible for ligation (Melamed et al., 2016).

For the RIL-seq experiments, the authors harvested the cells at 4°C and they resuspend the cells in ice-cold PBS prior to UV irradiation (Melamed et al., 2018, 2016), which results in a cold-shock that can affect the sRNA-interactome as well as sRNA stability. We cross-link actively growing cells in their growth medium and we UV irradiate our cells only for several seconds using the Vari-X-linker we recently developed (van Nues et al., 2017). We use filtration devices to rapidly harvest our cells (less than 30 seconds) and the filtered cells are subsequently stored at −80°C. We previously showed that filtration combined with short UV cross-linking times dramatically reduces noise introduced by the activation of the DNA damage response and significantly increased the recovery of short-lived RNA species (van Nues et al., 2017). We speculate that many of the interactions that are unique to our Hfq CLASH data represent short-lived RNA duplexes that are preferentially captured with our UV cross-linking and rapid cell filtration setup.

### The 3’UTR derived sRNA MdoR functions in a mixed feed forward loop by suppressing opposing pathways

We found a large number of chimeras that represent over a hundred distinct intermolecular interactions between 3’UTRs and other mRNA regions, which implicate direct mRNA-mRNA communication. These interactions could have been formed by 3’UTR fragments that have been processed by RNase E (or other RNases) or by an sRNA located within the 3’UTR that transcribed from an internal promoter. These 3’UTR fragments are primarily described as decoys or sponges for other sRNAs (Miyakoshi et al., 2015b), but could act as *trans*-acting sRNAs as well (Chao and Vogel, 2016). Among the interactions in our data we found 79 interactions between 3’UTRs and 5’UTRs (Supplementary Table 2.6). We speculate that many of these represent novel 3’UTR-derived sRNAs that target 5’UTRs of mRNAs. Reanalysis of RNA-seq data from a study that globally mapped *E. coli* transcription start sites (TSS) (Thomason et al., 2015) identified 47 TSS in these 3’UTR, strongly suggesting that these 3’UTRs host sRNAs. The majority of these 3’UTR fragments appear to have 5’ monophosphates. We also verified some of our findings by Northern blot analyses. Thus, we have potentially uncovered many novel 3’UTR-derived sRNAs.

Some of these predictions were validated by our follow up work (discussed below) and, while this work was in progress, also by others (Melamed et al., 2016). One of the 3’UTR-derived sRNAs we uncovered (MdoR) was of particular interest as it is only detected during the transition from late exponential to early stationary phase. A model of how MdoR contributes to maltose uptake is shown in Figure 9A.

The genetic structure and transcriptional regulation of the *mal* regulon are well understood. However, its post-transcriptional regulation has remained largely unexplored. Our work uncovered new links between the maltose uptake (*mal*) regulon, envelope stress-responses and membrane composition/assembly pathways. While initially cells metabolize more favourable carbon sources such as glucose, these are generally rapidly depleted and bacteria need to quickly switch to alternative carbon sources, such as maltose and maltodextrins. During maltose utilisation, the *malEFG* operon is transcribed by the MalT transcription factor and the transcript is processed by RNase E as well as other degradosome components (Khemici and Carpousis, 2004). Here we show that the MdoR sRNA is a product of *malEFG* processing, which is protected from further degradation by Hfq.

MdoR is unique in a sense that it not only a 3’UTR derived sRNA that targets multiple pathways, but also because it is part of a mixed coherent feed forward loop (FFL) that we predict promotes maltose uptake via LamB. Efficient uptake of maltose/maltodextrin not only requires the inner membrane transporters encoded by the *malEFG* operon, but also the high-affinity maltose transporter LamB (Figure 9A), which cooperates with the inner membrane proteins to import these carbon sources. *lamB* is significantly upregulated when cells start to utilize alternative carbon sources. A major advantage of a coherent FFL is that it can accelerate or delay responses to stimuli, such as changes in carbon source availability. In this FFL, the key activator is the MalT transcription factor, which induces the expression of both *malEFG* and *lamB* when cells start to consume maltose (Figure 9B). Expression of new OMPs, however, needs to be carefully coordinated as any changes in the protein composition of the OM, such as increased levels of LamB (or accumulation of misfolded LamB in the periplasm), can lead to induction of the σ^E^ envelope stress response caused by the reorganisation of the membrane when maltodextrins are utilised (Figure 9A)(Kenyon et al., 2005). We propose that *malEFG* indirectly promotes LamB accumulation via the MdoR sRNA that (a) reduces the levels of general porins (OmpC and OmpA) and (b), likely indirectly, dampens the σ^E^ stress response as a result of OMP level reduction. Dampening σ^E^ expression important during maltose uptake as it induces the expression of MicA, which directly downregulates *lamB* translation (Bossi and Figueroa-Bossi, 2007). When ectopically expressed at high levels, MdoR reduces mRNA levels of *ompC* and *ompA*, which may free up the resources enabling efficient accumulation of LamB in the outer membrane. At physiological levels MdoR significantly reduces MicA expression and we observed a ∼20% increase in *lamB* mRNA levels. *OmpC* mRNA levels were not substantially affected. While these changes appear modest, it is important to take into consideration the very high abundance and intrinsic stability of *lamB* and *ompC* mRNAs (minute-long half-lives). It is therefore expected that even a mild reduction in their mRNA levels can profoundly relieve the pressure on the OMP translation and assembly pathways (Guo et al., 2014). The net outcome of *mal* regulon transcription, MdoR biogenesis and regulatory activity, is increased expression of high-affinity components of maltose-specific transport (MalE and LamB).

MdoR is reminiscent of the *Vibrio cholerae* MicX sRNA that also mapped to the 3’ region of *malG* and regulates levels of outer membrane proteins (Davis and Waldor, 2007). However, the MicX targets are different from MdoR. Furthermore, unlike MdoR, MicX is produced from an independent promoter: Conditions that change the expression of the upstream Mal operon in *V. cholerae* do not affect accumulation of MicX (Davis and Waldor, 2007), whereas the levels of MdoR strongly correlate with *malEFG* mRNA levels. Nevertheless, both studies demonstrate that an sRNA resides in the 3’UTR of the *malG* transcript and pinpoint a mechanism where the expression of multiple porins is regulated by sRNAs encoded within transporter mRNAs.

### CLASH reveals large sRNA-sRNA interaction networks

Our analyses unearthed an unexpectedly large number (>100) of sRNA-sRNA interactions. The majority of these interactions involved known sRNA seed sequences, suggesting that these could represent *bona fide* interactions that prevent sRNAs from base-pairing with their targets or result in the degradation of the sRNAs. About 40 sRNA-sRNA interactions involving sRNAs from the core genome were detected in the RNase E CLASH dataset, about a quarter of which were also detected in our Hfq data. The relatively low overlap between the datasets is not surprising, given the differences in the growth conditions (virulence inducing conditions for enterohaemorrhagic *E. coli* vs growth transitions for commensal *E. coli* MG1655). Moreover, many of the sRNA-sRNA interactions recovered in association with RNase E are likely duplexes in the process of being degraded. Many of the sRNA-sRNA interactions unique to our Hfq data may represent interactions between anti-sRNAs and seed sequences of sRNAs which may not necessarily involve recruitment/activity of RNase E.

The chimeras containing CyaR fragments were of particular interest as CyaR is preferentially expressed during the transition from late exponential to stationary phase (De Lay and Gottesman, 2009; Wassarman et al., 2001) and may therefore play an important role in adaptation to nutrient availability. We could show that overexpression of ArcZ, which base-pairs with CyaR in our CLASH data, significantly reduced CyaR levels (Figure 9B). Using seed mutants and compensatory mutations, we confirmed direct interactions between ArcZ and CyaR and our data suggests that this interaction specifically takes place during the transition phase. Interestingly in *Salmonella*, overexpression of ArcZ showed a dramatic reduction in CyaR bound to Hfq and upregulation of CyaR targets, such as *nadE* (Papenfort et al., 2009), suggesting that this activity is conserved between these two Gram-negative bacteria. A similar type of asymmetric regulation has also been elegantly demonstrated for the Qrr3 sRNA of Vibrio cholera (Feng et al., 2015). The fate of these sRNA-sRNA duplexes may depend on the position of the interaction; It was shown that if sRNA-target RNA base-pair within a stabilizing 5’ stem, the sRNA will be preferentially degraded (Feng et al., 2015). Consistent with this, folding of the chimeric reads suggests that ArcZ preferentially base-pairs with CyaR at the 5’ end (Figure 4), which may alter secondary structures that normally help to stabilize the sRNA. The biological role of ArcZ targeting CyaR is unclear, however, a possible function could be to reduce noise in CyaR expression by preventing CyaR levels from overshooting during the transition phase.

ArcZ and CyaR target mRNAs are associated with many different processes. Thus, these interactions are expected to connect multiple pathways. For example, ArcZ regulation of CyaR may connect adaptation to stationary phase/biofilm development (De Lay and Gottesman, 2009; Monteiro et al., 2012) to quorum sensing and cellular adherence (De Lay and Gottesman, 2009). CyaR expression is controlled by the global regulator Crp. Most of the genes controlled by Crp are involved in transport and/or catabolism of amino acids or sugar. Interestingly, ArcZ downregulates the *sdaCB* dicistron which encodes for proteins involved in serine uptake and metabolism (Papenfort et al., 2009). This operon has been shown to be regulated by Crp as well, suggesting that ArcZ can counteract the activity of Crp.

## Materials and Methods

### Bacterial strains and culture conditions

An overview of the bacterial strains used in this study is provided in Supplementary Table 5. The *E. coli* MG1655, TOP10 or TOP10F’ strains served as parental strains. The *E. coli* K12 strain used for CLASH experiments, MG1655 *hfq*::HTF was previously reported (Tree et al., 2014). Cells were grown in Lysogeny Broth (LB) or minimal medium with supplements (1xM9 salts, 2 mM MgSO_4_,0.1 mM CaCl_2_, 0.03 mM thiamine, 0.2% carbon-source) at 37°C under aerobic conditions with shaking at 200 rpm. The media were supplemented with antibiotics where required at the following concentrations: ampicillin (Sigma, UK, A9518) – 100 µg/ml, chloramphenicol (Corning, US, C239RI) - 25 µg/ml, kanamycin (Gibco, US, 11815-024) - 50 µg/ml. Where indicated, 0.2% glucose or maltose were used. For induction of sRNA expression from plasmids, 1 mM IPTG, 200 nM anhydrotetracycline hydrochloride (Sigma, 1035708-25MG) or 0.2% L-arabinose (Sigma, A3256) were used.

### Construction of sRNA expression plasmids

For the pulse-overexpression constructs, the sRNA gene of interest was cloned at the transcriptional +1 site under P*ara* control by amplifying the pBAD+1 plasmid (Supplementary Table 5) by inverse PCR using Q5 DNA Polymerase (NEB). The pBAD+1 template is derived from pBAD*myc*HisA (Tree et al., 2014). The sRNA genes and seed mutants (SM) were synthesized as ultramers (IDT; Supplementary Table 5), which served as the forward primers, as described (Tree et al., 2014). The reverse primer (oligo pBAD+1_5P_rev) bears a monophosphorylated 5’-end to allow blunt-end self-ligation. The PCR reaction was digested with 10U DpnI (NEB) for 1h at 37°C and purified by ethanol precipitation. The sRNA-pBAD linear PCR product was circularized by self-ligation, performed as above. Ligations were transformed in DH5α competent cells. Positive transformants were screened by sequencing. The control plasmid pBAD+1 was constructed similarly by self-ligation of the PCR product generated from oligonucleotides pBAD+1_XbaI_fwd and pBAD+1_5P_rev. Small RNA overexpression constructs derived from the pZA21MCS and pZE12*luc* (Expressys) were generated identically, using the indicated ultramers in Supplementary Table 5 as forward primers, and oligos pZA21MCS_5P_rev and pZE12_5P_rev as reverse primers, respectively, and transformed in *E. coli* TOP10F’.

### Construction of mRNA-superfolder GFP fusions

Supplementary Table 5 lists all the plasmids, gene fragments and primers used for cloning procedures in this work. To construct constitutively expressed, in-frame mRNA-*sfGFP* fusions for the fluorescence reporter studies, the 5’UTR, start codon and first ∼5 codons of target genes were cloned under the control of PLtetO-1 promoter in a pXG10-SF backbone as previously described (Corcoran et al., 2012; Urban and Vogel, 2007). Derivatives of the target–GFP fusion plasmids harbouring seed mutations (SM) were generated using synthetic mutated gene-fragments (IDT, Belgium, Supplementary Table 5). To prepare the inserts, the target region of mRNA of interest was either amplified by PCR from *E. coli* genomic DNA or synthesized as g-blocks (IDT, Belgium) and cloned using NheI and NsiI restriction sites. Transformants were screened by restriction digest analysis and verified by Sanger sequencing (Edinburgh Genomics, Edinburgh, UK).

### Hfq UV Cross-linking, Ligation and Analysis of Hybrids (Hfq-CLASH)

CLASH was performed essentially as described(Waters et al., 2016), with a number of modifications including changes in incubation steps, cDNA library preparation, reaction volumes and UV cross-linking. *E. coli* expressing the chromosomal Hfq-HTF were grown overnight in LB at 37°C with shaking (200 rpm), diluted to starter OD_600_ 0.05 in fresh LB, and re-grown with shaking at 37°C in 750 ml LB. A volume of culture equivalent to 80 OD_600_ per ml was removed at the following cell-densities (OD_600_): 0.4, 0.8, 1.2, 1.8, 2.4, 3.0 and 4.0, and immediately subjected to UV (254 nm) irradiation for 22 seconds (∼500 mJ/cm2) in the Vari-X-linker (van Nues et al., 2017) (https://www.vari-x-link.com). Cells were harvested using a rapid filtration device (van Nues et al., 2017) (https://www.vari-x-link.com) onto 0.45 μM nitrocellulose filters (Sigma, UK, HAWP14250) and flash-frozen on the membrane in liquid nitrogen. Membranes were washed with ∼15 ml ice-cold phosphate-buffered saline (PBS), and cells were harvested by centrifugation. Cell pellets were lysed by bead-beating in 1 volume per weight TN150 buffer (50mM Tris pH 8.0, 150 mM NaCl, 0.1% NP-40, 5 mM β-mercaptoethanol) in the presence of protease inhibitors (Roche, A32965), and 3 volumes 0.1 mm Zirconia beads (Thistle Scientific, 11079101z), by performing 5 cycles of 1 minute vortexing followed by 1-minute incubation on ice. One additional volume of TN150 buffer was added. To reduce the viscosity of the lysate and remove contaminating DNA the lysate was incubated with RQ1 DNase I (10U/ml Promega, M6101) for 30 minutes on ice. Two-additional volumes of TN150 were added and mixed with the lysates by vortexing. The lysates were centrifuged for 20 minutes at 4000 rpm at 4°C and subsequently clarified by a second centrifugation step at 13.4 krpm, for 20 min at 4°C. Purification of the UV cross-linked Hfq-HTF-RNA complexes and cDNA library preparation was performed as described (Granneman et al., 2009). Cell lysates were incubated with 50 μl of pre-equilibrated M2 anti-FLAG beads (Sigma, M8823-5ML) for 1-2 hours at 4°C. The anti-FLAG beads were washed three times 10 minutes with 2 ml TN1000 (50 mM Tris pH 7.5, 0.1% NP-40, 1M NaCl) and three times 10 minutes with TN150 without protease inhibitors (50 mM Tris pH 7.5, 0.1% NP-40, 150mM NaCl). For TEV cleavage, the beads were resuspended in 250 μl of TN150 buffer (without protease inhibitors) and incubated with home-made GST-TEV protease at room temperature for 1.5 hours. The TEV eluates were then incubated with a fresh 1:100 dilution preparation of RNaceIt (RNase A and T1 mixture; Agilent, 400720) for exactly 5 minutes at 37°C, after which they were mixed with 0.4g GuHCl (6M, Sigma, G3272-100G), NaCl (300mM), and Imidazole (10mM, I202-25G). Note this needs to be carefully optimized to obtain high-quality cDNA libraries. The samples were then transferred to 50 μl Nickel-NTA agarose beads (Qiagen, 30210), equilibrated with wash buffer 1 (6 M GuHCl, 0.1% NP-40, 300 mM NaCl, 50 mM Tris pH 7.8, 10 mM Imidazole, 5 mM beta-mercaptoethanol). Binding was performed at 4°C overnight with rotation. The following day, the beads were transferred to Pierce SnapCap spin columns (Thermo Fisher, 69725), washed 3 times with wash buffer 1 and 3 times with 1xPNK buffer (10 mM MgCl_2_, 50mM Tris pH 7.8, 0.1% NP-40, 5 mM beta-mercaptoethanol). The washes were followed by on-column TSAP incubation (Thermosensitive alkaline phosphatase, Promega, M9910) treatment for 1h at 37°C with 8 U of phosphatase in 60 μl of 1xPNK, in the presence of 80U RNasin (Promega, N2115). The beads were washed once with 500 μl wash buffer 1 and three times with 500 μl 1xPNK buffer. To add 3’-linkers (App-PE – Supplementary Table 5), the Nickel-NTA beads were incubated in 80 μl 3’-linker ligation mix with (1 X PNK buffer, 1 µM 3’-adapter, 10% PEG8000, 30U Truncated T4 RNA ligase 2 K227Q (NEB, M0351L), 60U RNasin). The samples were incubated for 4 hours at 25°C. The 5’-ends of bound RNAs were radiolabelled with 30U T4 PNK (NEB, M0201L) and 3μl ^32^P-γATP (1.1µCi; Perkin Elmer, NEG502Z-500) in 1xPNK buffer for 40 min at 37°C, after which ATP (Roche, 11140965001) was added to a final concentration of 1mM, and the incubation prolonged for another 20 min to complete 5’-end phosphorylation. The resin was washed three times with 500 μl wash buffer 1 and three times with equal volume of 1xPNK buffer. For on-bead 5’-linker ligation, the beads were incubated 16h at 16°C in 1xPNK buffer with 40U T4 RNA ligase I (NEB, M0204L), and 1 μl 100 μM L5 adapter (Supplementary Table 5), in the presence of 1mM ATP and 60U RNasin. The Nickel-NTA beads were washed three times with wash buffer 1 and three times with buffer 2 (50 mM Tris–HCl pH 7.8, 50 mM NaCl, 10 mM imidazole, 0.1% NP-40, 5 mM β-mercaptoethanol). The protein-RNA complexes were eluted in two steps in new tubes with 200 μl of elution buffer (wash buffer 2 with 250 mM imidazole). The protein-RNA complexes were precipitated on ice by adding TCA (T0699-100ML) to a final concentration of 20%, followed by a 20-minute centrifugation at 4°C at 13.4 krpm. Pellets were washed with 800 μl acetone, and air dried for a few minutes in the hood. The protein pellet was resuspended and incubated at 65°C in 20 μl 1x NuPage loading buffer (Thermo Scientific, NP0007), resolved on 4–12% NuPAGE gels (Thermo Scientific, NP0323PK2) and visualised by autoradiography. The cross-linked proteins-RNA were cut directly from the gel and incubated with 160 μg of Proteinase K (Roche, 3115801001) in 600 μl wash buffer 2 supplemented with 1% SDS and 5 mM EDTA at 55°C for 2-3 hours with mixing. The RNA was subsequently extracted by phenol-chloroform extraction and ethanol precipitated. The RNA pellet was directly resuspended in RT buffer and was transcribed in a single reaction with the SuperScript IV system (Invitrogen, 18090010) according to manufacturer’s instructions using the PE_reverse oligo as primer. The cDNA was purified with the DNA Clean and Concentrator 5 kit (Zymo Research) and eluted in 11 μl DEPC water. Half of the cDNA (5 μl) was amplified by PCR using Pfu Polymerase (Promega, M7745) with the cycling conditions (95°C for 2 min; 20-24 cycles: 95°C for 20s, 52°C for 30s and 72°C for 1 min; final extension of 72°C for 5 min). The PCR primers are listed in Supplementary Table 5. PCR products were treated with 40U Exonuclease 1 (NEB, M0293L) for 1 h at 37°C to remove free oligonucleotide and purified by ethanol precipitation/ or the DNA Clean and Concentrator 5 kit (Zymo Research, D4003T). Libraries were resolved on a 2% MetaPhor agarose (Lonza, LZ50181) gel and 175-300bp fragments were gel-extracted with the MinElute kit (Qiagen, 28004) according to manufacturer’s instructions. All libraries were quantified on a 2100 Bionalyzer using the High-Sensitivity DNA assay and a Qubit 4 (Thermo Scientific, Q33226). Individual libraries were pooled based on concentration and barcode sequence identity. Paired-end sequencing (75 bp) was performed by Edinburgh Genomics on an Illumina HiSeq 4000 platform.

### RNA-seq

*E. coli* MG1655 was cultured, UV-irradiated and harvested as described for the CLASH procedure. Total RNA was extracted using the Guanidium thiocyanate phenol method. RNA integrity was assessed with the Prokaryote Total RNA Nano assay on a 2100 Bioanalyzer (Agilent, G2939BA). Sequencing libraries from two biological replicates were prepared by NovoGene using the TruSeq library preparation protocol and 150bp paired-end sequencing was performed on an Illumina NovaSeq 6000 system. This yielded ∼7-8 million paired-end reads per sample. For the overexpression analysis of MdoR, we generated RNA-seq libraries using an in-house protocol. Genomic DNA was removed by incubating 10 μg of total RNA with 2U Turbo DNase (Thermo Scientific, AM2238) in a 50 μl final volume for 30 minutes at 37°C in the presence of 10 U SuperaseIn RNase Inhibitor (Thermo Scientific, AM2694). RNA was subsequently phenol-chloroform extracted and purified by ethanol-precipitation. Ribosomal RNA was removed with the Ribo-Zero rRNA Removal Kit (Gram-Negative Bacteria; Illumina, MRZGN126) according to the manufacturer’s instructions. Successful rRNA depletion was verified on the Agilent 2100 Bioanalyzer. The RNA was fragmented for 5 min at 95°C in the presence of Superscript III buffer (Invitrogen) followed by a five-minute incubation on ice. Reverse-transcription (RT) was performed with Superscript III (Thermo Scientific, 18080044) in 20 μl reactions according to manufacturer’s procedures using 250 ng of ribosomal RNA depleted RNA and 2.5 μM random hexamers (PE_solexa_hexamer, oligo 73, Supplementary Table 5). The RNA and free primers were degraded using 20U of Exonuclease I (NEB, M0293L) and 50U RNaseIf (NEB, M0243S) and the cDNA was purified with the DNA Clean & Concentrator 5 kit (Zymo Research). Ligation of the 5’ adapter (P5_phospho_adapter, oligo 39) to the cDNA was performed using CircLigase II (Lucigen, CL9021K) for 6 hours at 60°C, followed by a 10-minute inactivation at 80°C. The cDNA was purified with the DNA Clean & Concentrator 5 kit. Half of the cDNA library was PCR amplified using Pfu polymerase (Promega, M7745) using the P5 forward PCR oligonucleotide and barcoded BC reverse oligonucleotides (200 nM; Supplementary Table 5; 95°C for 2 min, 95°C for 20s, 52°C for 30s and 72°C for 1 min, and a final extension of 72°C for 5 min. 20 cycles of amplification). The PCR products were treated with Exonuclease 1 (NEB, M0293L) for 1 h at 37°C and purified by ethanol precipitation. Libraries were resolved on a 2% MetaPhor agarose gel 200-500 bp fragments were gel-extracted using the MinElute kit. All libraries were quantified on a 2100 Bionalyzer using the High-Sensitivity DNA assay (Agilent, 5067-4627). Individual libraries were pooled in equimolar amounts. Paired-end sequencing (75 bp) was performed by Edinburgh Genomics on the Illumina HiSeq 4000 platform.

### Small RNA overexpression studies

Individual TOP10F’ clones carrying pZA21 and pZE12-derived sRNA constructs and control plasmids combinations (Supplementary Table 5) were cultured to OD_600_ 0.1 and expression of sRNAs was induced with IPTG and anhydrotetracycline hydrochloride (Sigma, I6758-1G and 1035708-25MG) for one hour. Cells were collected by centrifugation for 30 seconds at 14000 rpm, flash-frozen in liquid nitrogen and total RNA was isolated as above. Genomic DNA was digested with Turbo DNase (Thermo Scientific, AM2238), then the RNA was purified with RNAClean XP beads (Beckman Coulter, A63987). Gene expression was quantified by RT-qPCR (see below) using 10 ng total RNA as template, and expressed as fold change relative to the reference sample containing pJV300 (Sittka et al., 2007) or pZA21.

For pulse-overexpression studies overnight MG1655 cultures containing pBAD::sRNA and empty pBAD+1 control plasmids were inoculated in fresh LB-ampicillin medium at a starting OD_600_ of 0.05 and grown aerobically at 37°C to OD_600_ 0.4. Pre-induction (0 min) and post-induction samples were harvested. For induction, cultures were supplemented with L-arabinose (Sigma, A3256) and rapidly collected by filtration and flash-frozen in liquid nitrogen at the indicated time-points. RNA was extracted from three biological replicate time-series, followed by RNA-seq library preparation, next generation sequencing and DESeq2 analysis of differentially expressed genes.

### GFP reporter system to quantify sRNA effect on target expression

A two-plasmid system was used to express each sRNA, and mRNA-*sfGFP* fusions (Corcoran et al., 2012; Urban and Vogel, 2007) with modifications. The sRNA and sfGFP-fusion plasmids were co-transformed in *E. coli* TOP10 cells by electroporation and cells were maintained on dual selection with ampicillin and chloramphenicol. In TOP10 cells, the mRNA-*sfGFP* constructs are constitutively expressed, whereas sRNA expression requires L-arabinose induction. The expression of sfGFP-fused targets in the presence or absence of sRNAs was quantified at the protein level, by plate reader experiments and at the RNA level, by RT-qPCR.

For the plate reader experiments, a single colony of bacterial strain harbouring a sRNA-target-sfGFP combination was inoculated in a 96-well Flat Bottom Transparent Polystyrene plate with lid (Thermo Scientific, 152038) and cultured overnight at 37°C in 100 μl LB supplemented with antibiotics and L-arabinose (Sigma, A3256) to induce expression of sRNAs. Next day, each overnight inoculum was diluted 1:100 by serial dilution, in triplicate, in LB with freshly prepared L-arabinose to a final volume of 100 μl. Cultures were grown in a 96-well plate in an Infinite 200 Pro plate reader (Tecan) controlled by i-control software (Tecan) for 192 cycles at 37°C with 1 min orbital shaking (4 mm amplitude) every 5th minute. To monitor optical density over time, the following parameters were used: wavelength 600 nm, bandwidth 9 nm. Fluorescence was monitored with excitation wavelength 480 nm, bandwidth 9 nm and emission wavelength 520 nm, bandwidth 20 nm. Measurements were recorded at 5-minute intervals, by top reading. Raw data was processed following guidance from previous reports (Urban and Vogel, 2007). First, the range of linearity of increase of fluorescence with OD_600_ was identified for all individual triplicates. Only the linearity range common to all triplicates was considered for further analysis. For each set of triplicates, the mean fluorescence was calculated at each OD_600_. To correct for background and cell autofluorescence, the mean fluorescence of a strain with plasmid pXG-0 was subtracted from all strains with GFP plasmids at the equivalent OD_600_. Ultimately, a curve was generated for each sample, plotting the background-corrected fluorescence (GFP) versus OD_600_. The experiments were performed for three biological replicates, and mean values and standard error of the means calculated for each strain.

### RT-qPCR

Total RNA (12.5µg) was treated with 2U of Turbo DNase (Thermo Scientific, AM2238) for 1 hour at 37°C in a 10 μl reaction in the presence of 2U SuperaseIn RNase inhibitor (Thermo Scientific, AM2694). The DNase was inactivated by 10 minutes incubation at 75°C. Reverse transcription (RT) was performed in a single reaction for all target genes of interest using a mix of gene-specific RT primers at 3.5 μM concentration each. After addition of 2.5 μl RT primer mix, the RNA and primers were denatured at 70°C for 3 min, then snap chilled and incubated on ice for 5 min. RT was performed for 1 hour at 55°C with SuperScript III (Thermo Scientific, 18080051) using 5 μl of RNA-RT primers mix in 10 μl final volume (100 U Superscript III, 2.5 mM DTT, 1xFS Buffer, 0.75 mM dNTPs) in the presence of 1U RNasin (Promega, N2115). RT was followed by treatment with 5U RNase H for 30 min at 37°C to remove the RNA from the RNA-cDNA duplexes. The cDNA was diluted 10-fold with DEPC water. Quantitative PCR was performed on 50ng of DNAse I-treated total RNA using the Brilliant III UltraFast SYBR Green QPCR Master Mix (Agilent, #600883) and the Luna Universal One-Step RT-qPCR Kit (NEB, E3005E) according to manufacturer’s instructions. The qPCRs were run on a LightCycler 480 (Roche), and the specificity of the product was assessed by generating melting curves, as follows: 65°C-60s, 95°C (0.11 ramp rate with 5 acquisitions per °C, continuous). The data analyses were performed with the IDEAS2.0 software, at default settings: Absolute Quantification/Fit Points for Cp determination and Melt Curve Genotyping.

The qPCR efficiency of primer pairs was assessed by performing standard curves by serial dilution of template RNA or genomic DNA. Negative controls such as -RT or no template control were used throughout, and the qPCR for all samples was performed in technical triplicate. Outliers from the samples with technical triplicate standard deviations of Cp > 0.3 were discarded from the analyses. To calculate the fold-change relative to the control, the 2^-ddCp^ method was employed, using *recA* or 5S rRNA (*rrfD*) as the reference genes where indicated. Experiments were performed for minimum two biological replicates, and the mean fold-change and standard error of the mean were computed. Unless otherwise stated, significance of the fold-change difference compared to the reference sample control (for which fold-change =1) was tested with a one-sample t-test.

### Northern Blot analysis

Total RNA was extracted from cell lysates by GTC-Phenol extraction. For large RNA fragments, 10 μg of total RNA was resolved on a 1.25% BPTE-gel (pH 7) and transferred to a nylon membrane (HyBond N+, GEHealthcare, RPN1210B) by capillarity. For short RNA fragments, 10 μg total RNA was separated on an 8% polyacrylamide TBE-Urea gel and transferred to a nitrocellulose membrane by electroblotting for four hours at 50 V. Membranes were pre-hybridised in 10 ml of UltraHyb Oligo Hyb (Thermo Scientific, AM8663) for one hour and probed with ^32^P-labeled DNA oligo at 42°C for 12-18 hours in a hybridization oven. The sequences of the probes used for Northern blot detection are detailed in Supplementary Table 5. Membranes were washed twice with 2xSSC + 0.5% SDS solution for 10 minutes and visualized using a Phosphor imaging screen and FujiFilm FLA-5100 Scanner (IP-S mode). For detection of highly abundant species (5S rRNA) autoradiography was used for exposure.

### Western blot analyses

A fraction of the *E. coli* MG1655 Hfq::*htf* lysates used for the RNA-seq experiments using strains cultured, cross-linked, harvested and lysed in identical conditions as the CLASH experiments containing 40 µg protein was run on PAGE gels and transferred to a nitrocellulose membrane. The membranes were blocked for one hour in blocking solution (5% non-fat milk in PBST (1X phosphate saline buffer, 0.1% Tween-20). To detect Hfq-HTF protein, the membrane was probed overnight at 4°C with the Rabbit anti-TAP polyclonal primary antibody (Thermo Fisher, 1:5000 dilution in blocking solution), which recognizes an epitope at the region between the TEV-cleavage site and His6. For the loading control we used a rabbit polyclonal to GroEL primary antibody (Abcam, 1:150000 dilution, ab82592), for 2 hours at room temperature. After 3×10 min PBST washes, the membranes were blotted for one hour with a Goat anti-rabbit IgG H&L (IRDye 800) secondary antibody (Abcam, ab216773, 1:10000 in blocking solution) at room temperature. Finally, after three 10-minute PBST washes, the blot was rinsed in PBS, and the proteins were visualised with a LI-COR (Odyssey CLx) using the 800 nm channel and scan intensity 4. Image acquisition and quantifications were performed with the Image Studio Software.

### Primer extension analysis

One microgram total RNA was reverse-transcribed using SuperScript III reverse transcriptase (Thermo Scientific, 18080051) using ^32^P-radiolabelled oligonucleotides as primers (Supplementary Table 5). Primers were added to the RNA and annealing was performed by heating the samples at 85°C for three minutes and then snap chilling them on ice. The RT was performed for one hour at 45°C, followed by Exonuclease I and RNaseIf (NEB M0293L and M0243S) (0.5 μl each) treatment for 30 minutes at 37°C. Reactions were stopped by mixing with an equal volume of 2XRNA loading dye (NEB, B0363S), 2 minutes incubation at 95°C and snap chilled. The sequencing ladders were prepared with Sequenase v2.0 (Thermo Scientific, 70775Y200UN) according to specified instructions. Samples were resolved on 6% PAA/8M TBE-urea gels and visualized using the FujiFilm FLA5100 scanner.

### Construction of the MdoR seed-mutant strain

To mutate the chromosomal copy of MdoR, we used the λRed system (Datsenko and Wanner, 2000). We amplified the integration cassette from plasmid pKD4 with ultramers 895 and 896, containing homology regions to the coding sequence of *malG*, the desired MdoR sequence and to the region immediately downstream of the Rho-independent terminator, respectively. With this design, the scar after removal of the *Kan*^r^ cassette was expected at a site outside the MdoR/*malG* sequence. The PCR product was electroporated in *E. coli* MG1566 strains carrying the pKD46 plasmid from which λRed recombinase was induced with 10 mM L-arabinose. Correct replacement of the MdoR seed sequence was screened by colony PCR using primer pairs: 725 & 909 and 726 & 910 (Supplementary Table 5). The antibiotic resistance cassette was removed from substitution mutants by FLP-recombinase expressed constitutively from pE-FLP (St-Pierre et al., 2013). Successful allele replacement was confirmed by Sanger sequencing.

### Polysome profiling analyses

Wild-type *E. coli* MG1655 containing empty pBAD plasmid and an isogenic strain containing pBAD::MdoR were grown in LB until OD_600_ 0.4, then treated for 15 minutes with L-arabinose (Sigma, A3256) to induce overexpression of MdoR, and cycloheximide (Sigma, C4859-1ML) at a final concentration of 100 µg/ml for 3 minutes. 200 ml of cells were harvested by rapid filtration and flash frozen. The cells were washed in ice-cold PBS supplemented with 100µg/ml cycloheximide.

Polysomal profiling was performed according to previously described protocols (Bernabò et al., 2017; Lunelli et al., 2016) with minor changes in the lysis buffer (10 mM NaCl, 10 mM MgCl_2_, 10 mM Tris-HCl (pH 7.5), 100 µg/ml cycloheximide, 1% (w/v) Na-Deoxycholate, 1U RQ DNAse I (Promega, M6101), 0.6U/mL RiboLock (Thermo Scientific, EO0381) in DEPC water). Lysates were kept on ice for 30 min, centrifuged 3X at 15000 g for 10 min. The supernatants were loaded on a linear 10%–30% [w/v] sucrose gradient and centrifuged for 4 hours using a SW41 rotor at 40000 rpm in a Beckman Optima XPN-100 Ultracentrifuge. Fractions of 1 mL in volume were collected monitoring the absorbance at 254 nm with the UA-6 UV/VIS detector (Teledyne Isco). Fractions from the entire gradient (total RNA) and from the fractions corresponding to ribosomes (70S) and polysomes (polysomal RNA) were pooled and RNA was purified by acid phenol–chloroform extraction according to (Tebaldi et al., 2012).

### Terminator™ 5′-PhosphateDependent Exonuclease treatment

Ten micrograms of total RNA extracted from cell-samples at OD_600_ 1.2 and 1.8 were treated with 5’-Terminator Dependent Exonuclease (Lucigen, TER51020) as per manufacturer instructions using Buffer A. The reaction was terminated by phenol extraction and ethanol precipitation, and the RNA was loaded on 8% polyacrylamide-urea gels and transferred to nylon membranes that were probed for MdoR, CpxQ, RybB and 5S rRNA (Supplementary Table 5).

### Seed mutant studies

Wild-type MG1655 and seed mutant strains were grown overnight in minimal medium with glucose. Next day, each starter culture was split and inoculated at OD_600_ 0.05 in fresh M9 medium with glucose or maltose as the sole carbon source. Growth was monitored and cells were harvested at OD_600_ 0.5. Total RNA was extracted, and gene expression was quantified by RT-qPCR or Northern Blot.

### Computational analysis

#### Pre-processing of the raw sequencing data

Raw sequencing reads in fastq files were processed using a pipeline developed by Sander Granneman, which uses tools from the pyCRAC package (Webb et al., 2014). The entire pipeline is available at https://bitbucket.org/sgrann/). The CRAC_pipeline_PE.py pipeline first demultiplexes the data using pyBarcodeFilter.py and the in-read barcode sequences found in the L5 5’ adapters. Flexbar then trims the reads to remove 3’-adapter sequences and poor-quality nucleotides (Phred score <23). Using the random nucleotide information present in the L5 5’-adaptor sequences, the reads are then collapsed to remove potential PCR duplicates. The reads were then mapped to the *E. coli* MG1655 genome using Novoalign (www.novocraft.com). To determine to which genes the reads mapped to, we generated an annotation file in the Gene Transfer Format (GTF). This file contains the start and end positions of each gene on the chromosome as well as what genomic features (i.e. sRNA, protein-coding, tRNA) it belongs to. To generate this file, we used the Rockhopper software (Tjaden, 2015) on *E. coli* rRNA-depleted total RNA-seq data (generated by Christel Sirocchi), a minimal GTF file obtained from ENSEMBL (without UTR information). The resulting GTF file contained information not only on the coding sequences, but also complete 5’ and 3’ UTR coordinates. We then used pyReadCounters.py with Novoalign output files as input and the GTF annotation file to count the total number of unique cDNAs that mapped to each gene.

#### Normalization steps

To normalise the read count data generated with pyReadCounters.py and to correct for differences in library depth between time-points, we calculated Transcripts Per Million reads (TPM) for each gene. Briefly, for each time-point the raw counts for each gene was first divided by the gene length and then divided by the sum of all the values for the genes in that time-point to normalize for differences in library depth. The TPM values for each OD_600_ studied were divided by the TPM values of the first sample (OD_600_ 0.4) and were then log_2_-normalized. The log_2_-normalized fold-changes were used to compare RNA-seq and Hfq-cross-linking profiles among samples, and to perform k-means clustering with the python sklearn. cluster.KMeans class.

#### Hfq-binding coverage plots

For the analysis of the Hfq binding sites the pyCRAC package (Webb et al., 2014) was used (versions. 1.3.2-1.4.4). The pyBinCollector tool was used to generate Hfq cross-linking distribution plots over genomic features. First, PyCalculateFDRs.py was used to identify the significantly enriched Hfq-binding peaks (minimum 10 reads, minimum 20 nucleotide intervals). Next, pyBinCollector was used to normalize gene lengths by dividing their sequences into 100 bins and calculate nucleotide densities for each bin. To generate the distribution profile for all genes individually, we normalized the total number of read clusters (assemblies of overlapping cDNA sequences) covering each nucleotide position by the total number of clusters that cover the gene. Motif searches were performed with pyMotif.py using the significantly enriched Hfq-binding peaks (FDR intervals). The 4-8 nucleotide k-mers with Z-scores above the indicated threshold were used for making the motif logo with the k-mer probability logo tool (Wu and Bartel, 2017) with the -ranked option (http://kplogo.wi.mit.edu/).

#### Analysis of chimeric reads

Chimeric reads were identified using the hyb package using default settings (Travis et al., 2013) and further analysed using the pyCRAC package (Webb et al., 2014). To apply this single-end specific pipeline to our paired-end sequencing data, we joined forward and reverse reads using FLASH (Magoč and Salzberg, 2011), which merges overlapping paired reads into a single read. These, as well as any remaining single reads, were then analysed using hyb. The -anti option for the hyb pipeline was used to be able to use a genomic *E. coli* hyb database, rather than a transcript database. Uniquely annotated hybrids (.ua.hyb) were used in subsequent analyses. To visualise the hybrids in the genome browser, the .ua.hyb output files were converted to the GTF format. To generate distribution plots for the genes to which the chimeric reads mapped, the parts of the chimeras were clustered with pyClusterReads.py and BEDtools (Quinlan and Hall, 2010) (intersectBed) was used to remove clusters that map to multiple regions. To produce the coverage plots with pyBinCollector, each cluster was counted only once, and the number of reads belonging to each cluster was ignored.

#### sRNA density plots

To visualize the nucleotide read density of sRNA-target pairs for a given sRNA, we first merged the hyb datasets for all OD_600_ and biological replicates and filtered the interactions that were found statistically significant in the unified dataset. For each sRNA-target pair in the filtered dataset, the hit counts at each nucleotide position for all chimeras were summed. The count data was log_2_-normalised (actually log_2_(Chimera count +1) to avoid NaN for nucleotide positions with 0 hits when log-transforming the data).

#### sRNA-sRNA network visualization

Only the sRNA-sRNA chimeric reads representing statistically significant interactions in the merged CLASH dataset were considered. For each such interaction, chimera counts corresponding in either orientation were summed, log_2_-transformed and visualized with the igraph Python package.

#### Differential expression analyses

For the differential expression analyses DESeq2 was used (Love et al., 2014). Three MdoR pulse-overexpression datasets were compared to three pBAD Control overexpression datasets. Only differentially expressed genes that had an adjusted p-value of 0.05 or lower were considered significant.

#### Multiple sequence alignments and conservation analyses

The homologous sequences of MdoR in other enterobacteria were retrieved by BLAST. JalView was used for the multiple sequence alignments, using the MAFFT algorithm (Waterhouse et al., 2009).

### Data and Code availability

The next generation sequencing data have been deposited on the NCBI Gene Expression Omnibus (GEO) with accession number GSE123050. The python pyCRAC(Webb et al., 2014), kinetic-CRAC and GenomeBrowser software packages used for analysing the data are available from https://bitbucket.org/sgrann (pyCRAC up to version 1.3.3), https://git.ecdf.ed.ac.uk/sgrannem/ and pypi (https://pypi.org/user/g_ronimo/). The hyb pipeline for identifying chimeric reads is available from https://github.com/gkudla/hyb. The FLASH algorithm for merging paired reads is available from https://github.com/dstreett/FLASH2. Bedgraph and Gene Transfer Format (GTF) generated from the analysis of the Hfq CLASH, RNA-seq and TEX RNA-seq data(Thomason et al., 2015) are available from the Granneman lab DataShare repository (https://datashare.is.ed.ac.uk/handle/10283/2915).

## Acknowledgement

We are grateful to Lionello Bossi and Meriem El Karoui for their valuable feedback on the project and fruitful discussions. We thank Christel Sirocchi for help with preparing *E. coli* RNA-seq libraries, Erica de Leau for expert technical assistance and the members of the Granneman lab for critically reading the manuscript. We thank Jean-Marie Pages and Julia Vergalli for help the analyses of LamB levels and for fruitful discussions. This work was supported by grants from the Wellcome Trust (091549 to S.G and 102334 to I.A.I.), the Wellcome Trust Centre for Cell Biology core grant (092076), a Medical Research Council non Clinical Senior Research Fellowship (MR/R008205/1 to S.G.), the Australian National Health and Medical Research Council Project grants (GNT1067241 and GNT1139313 to J.J.T) and the Autonomous Province of Trento (Axonomix to G.V and M.M.). Sequencing of Hfq CLASH data was carried out by Edinburgh Genomics that is partly supported through core grants from NERC (R8/H10/56), MRC (MR/K001744/1) and BBSRC (BB/J004243/1).

## Materials & Correspondence

All requests for code, materials and reagents should be sent to Sander Granneman

## Supplementary Figures and legends

**Figure 1–figure supplement 1.**
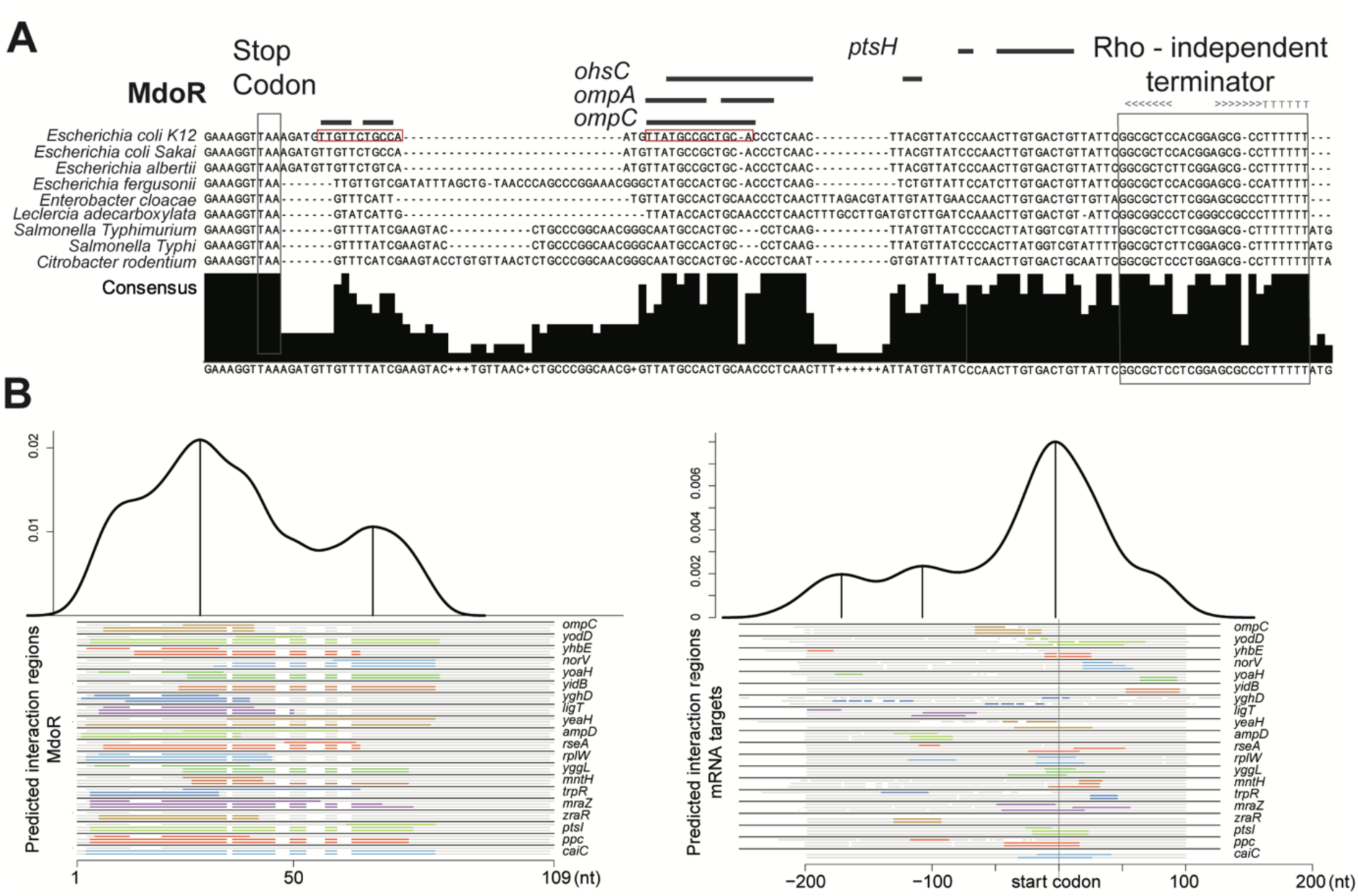
Hfq expression and Hfq binding to RNAs at different cell densities in UV-irradiated *E. coli*. **(A)** Western blot analyses of Hfq levels during various growth stages. Hfq-HTF was detected using an anti-TAP primary antibody, and a fluorescent secondary antibody. GroEL was used as a loading control. **(B)** Quantification of Hfq levels from the Western blot result. The fluorescent signal for Hfq-HTF and GroEL was measured with the LI-COR from biological replicate experiments. The levels of Hfq were normalised to GroEL and expressed as fold-change relative to OD_600_ 0.4. **(C)** Hfq crosslinking to RNA is similar at each optical density. Autoradiogram showing the purified radioactively labelled Hfq-RNA complexes for each OD_600_ after elution from the nickel beads. Source data for (**A**-**C**) are provided as a Source Data file.

**Figure 1–figure supplement 2.**
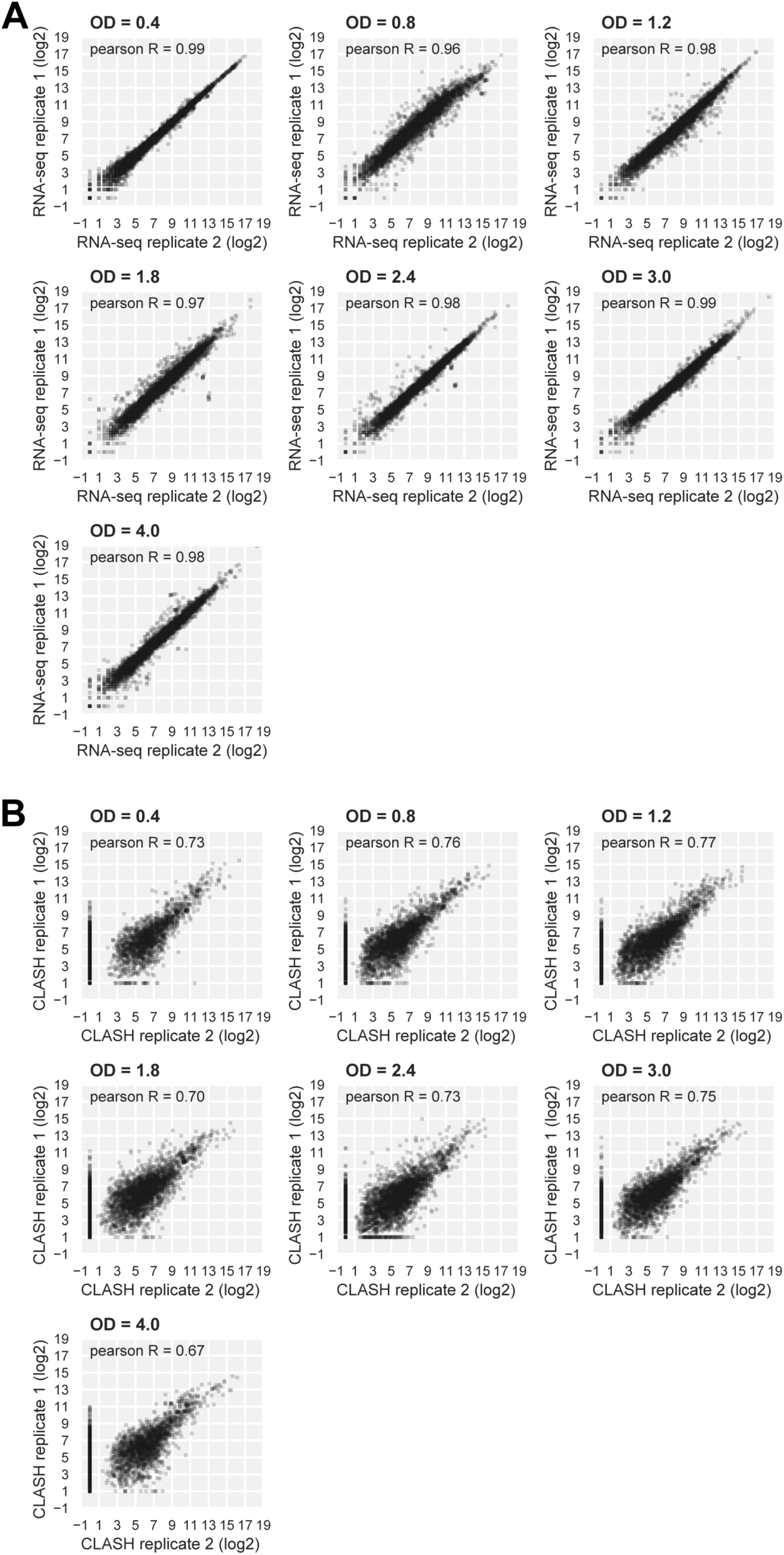
RNAseq and Hfq CLASH replicate datasets are highly correlated. (**A, B**) Scatter plots showing the distribution of log_2_ Transcripts Per Million (TPM) normalised read counts for Hfq CLASH (**A**) and RNA-seq (**B**) biological replicates. Pearson R coefficients describing the correlation between the two independent experiments at each OD_600_ are included.

**Figure 1–figure supplement 3.**
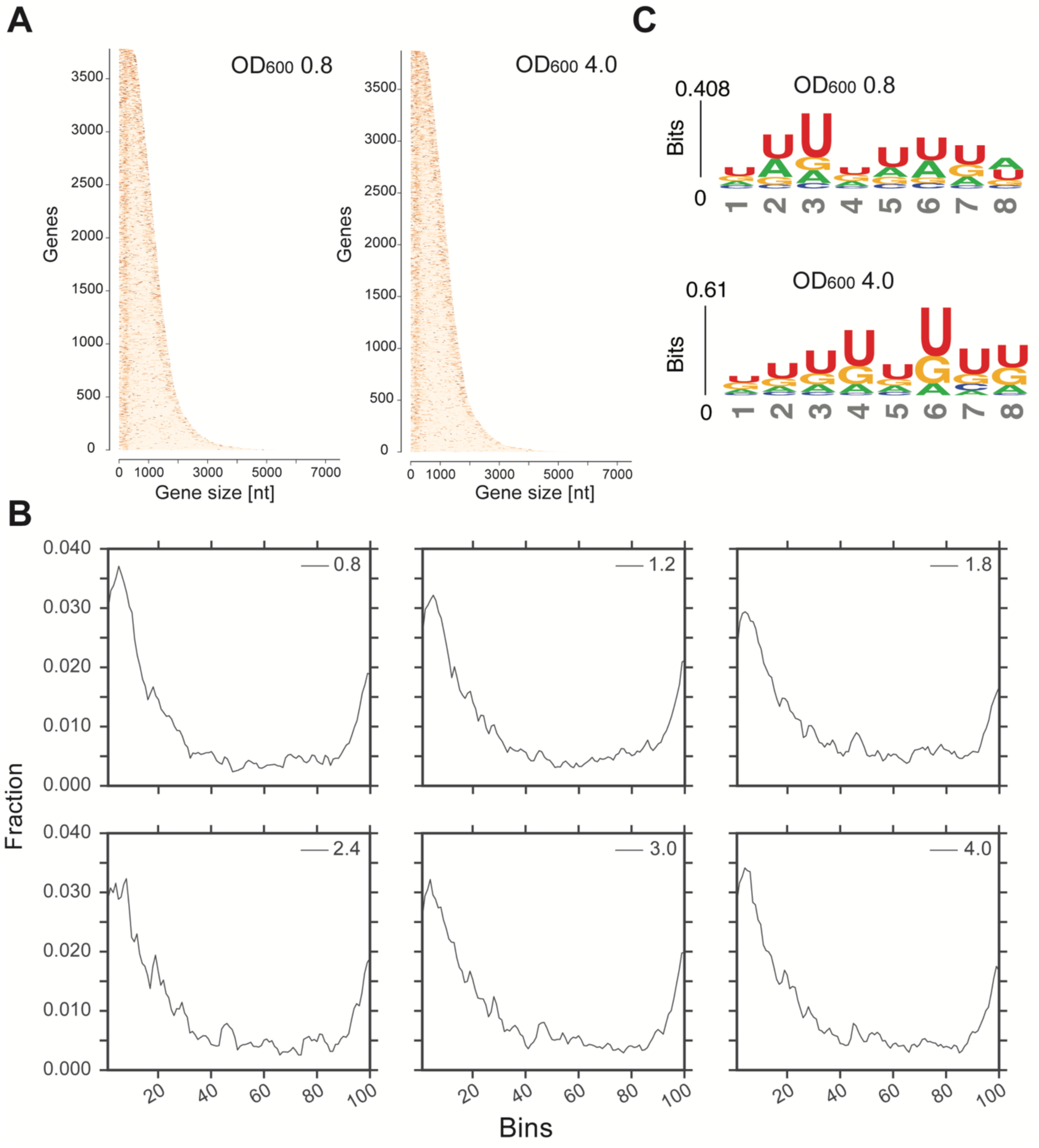
Transcriptome-wide maps of Hfq binding to mRNA genes. **(A)** Heatmaps showing the distribution of Hfq binding sites across all mRNA genes at OD_600_ 0.8 and 4.0. The genes are sorted by their sequence length (x-axis); the darker a nucleotide is, the more Hfq is crosslinked to it. To generate the heatmap, Hfq binding clusters were generated. A 5’-and 3’UTR length of 200bp was used. **(B)** Hfq binds to poly-U tracks. Significant k-mers (4-8 nt in length) were identified using the pyMotif tool of the pyCRAC package(Webb et al., 2014) and the motif logo was generated using all k-mers with a Z-score > 3, with kpLogo(Wu and Bartel, 2017). (**C**) A more stringent selection of the genes used to generate the distribution of Hfq binding to the transcriptome: all genes with overlapping 5’ or 3’UTRs were removed from the analysis to avoid ‘duplicate’ counting. For all remaining cDNAs, FDR intervals of minimum 20 nt were considered for distribution plotting. The interval length (with UTR flanks as in the GTF annotation file) for each gene was normalized over 100 bins (x-axis), and the fraction of hits in each bin was calculated (y-axis).

**Figure 2–figure supplement 1.**
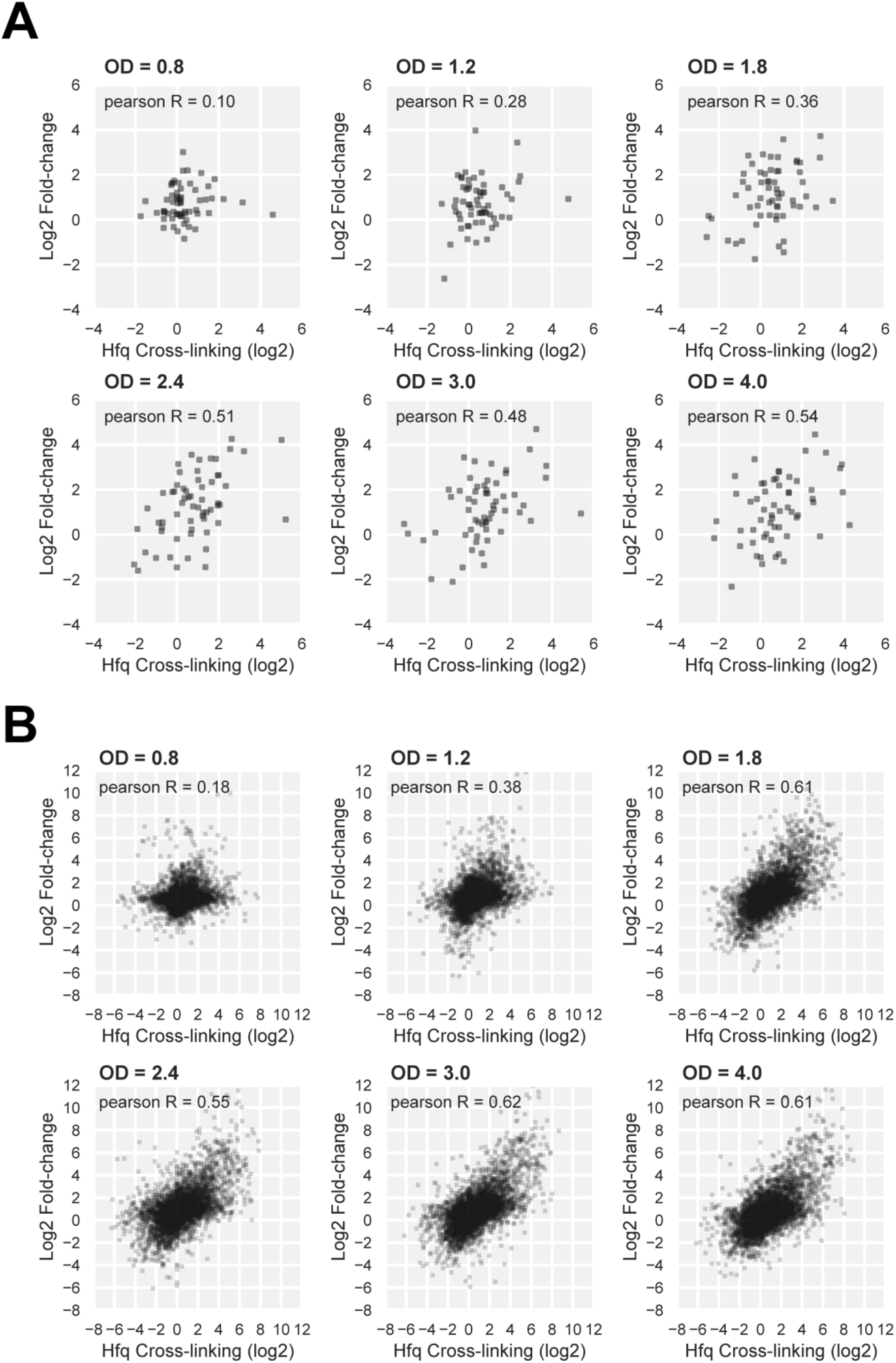
The sRNA and mRNA levels are not always strongly correlated with Hfq binding during the various stages of growth. Scatter plots comparing changes in Hfq binding (x-axis) to RNA levels (y-axis) for the indicated OD_600_ for sRNAs (**A**) and all transcript classes (**B**) show that the correlation improves at higher OD_600_. For each OD_600_, the Transcripts Per Million (TPM) normalised read counts at each density-point were divided by the OD_600_ 0.4 data, and the resulting ratios were log_2_-normalized. Pearson R coefficients describing the correlation between the two independent experiments at each OD_600_ is also included for each plot.

**Figure 4–figure supplement 1.**
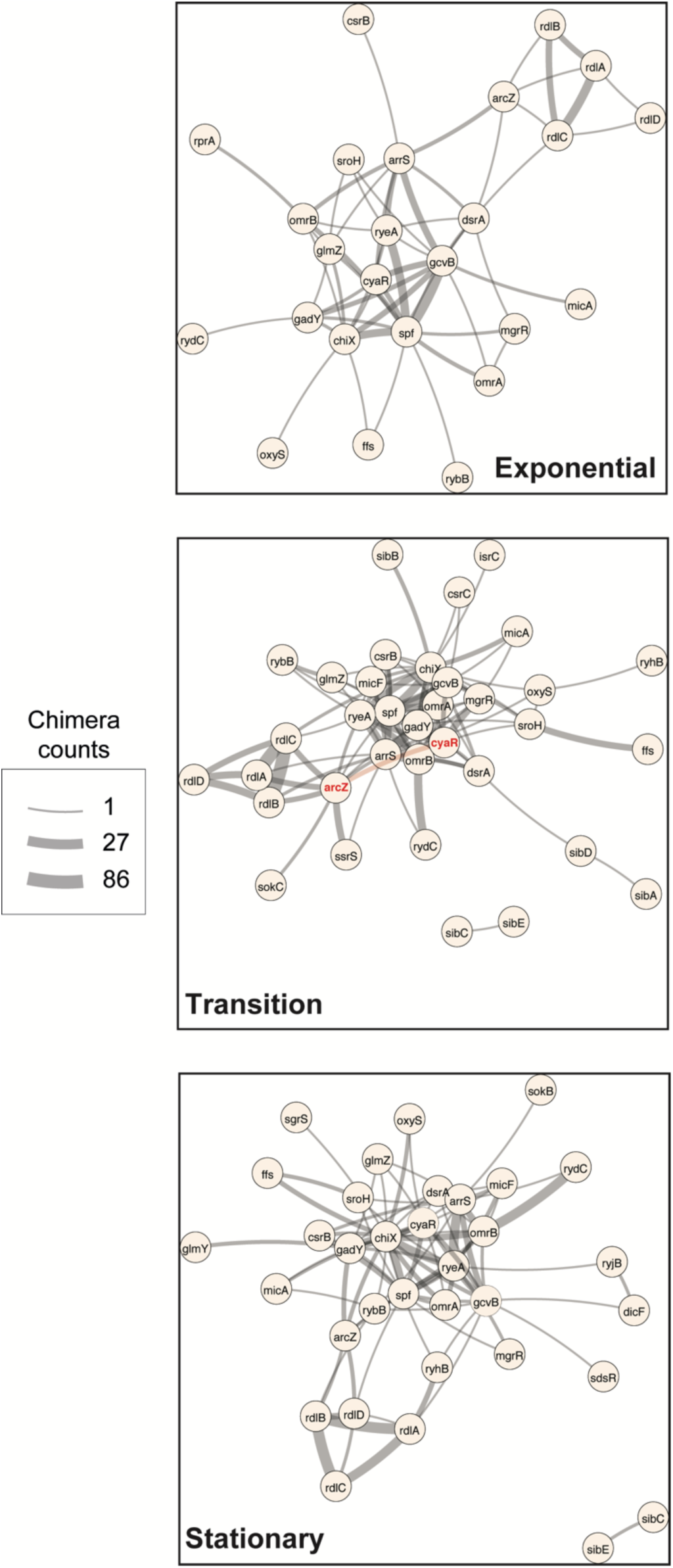
sRNA-RNA interactions identified by CLASH are growth-stage specific. sRNA-sRNA network generated from the statistically significant CLASH interactions from two biological replicates, recovered at three main growth stages: exponential (OD_600_ 0.4 and 0.8), transition (OD_600_ 1.2, 1.8, 2.4) and early stationary (OD_600_ 3.0 and 4.0). The thickness of the edges is proportional to the log_2_(unique chimera count for each interaction). Only sRNAs transcribed from independent promoters were included in the analysis.

**Figure 4–figure supplement 2.**
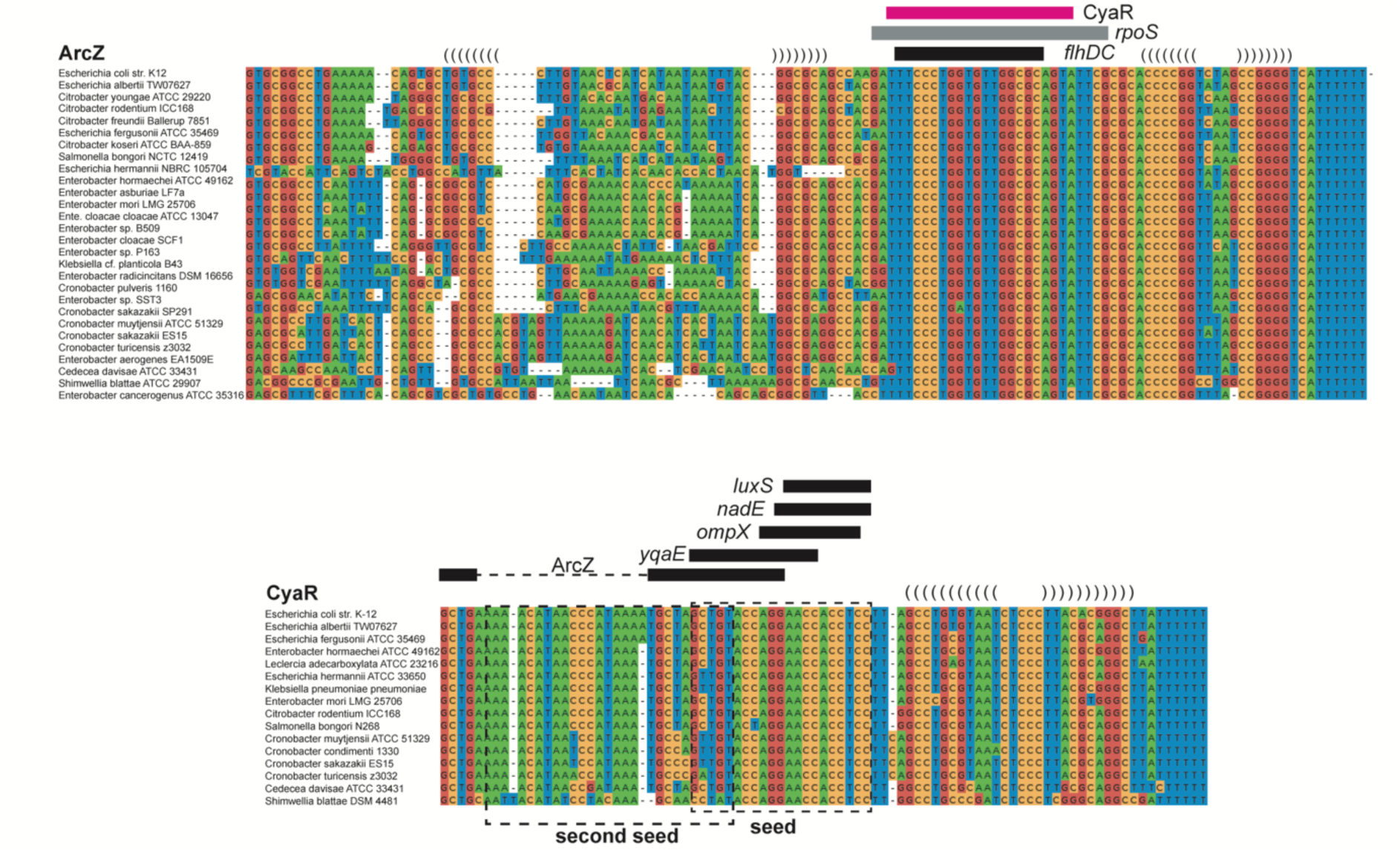
Interactions between ArcZ, CyaR and GcvB are conserved. Alignments of ArcZ, CyaR and GcvB were compiled as previously described(van Nues et al., 2016). Names of the enteric bacteria from which the sequence was retrieved are given on the left. Indicated are possible stem-loops (brackets), seed regions (boxed in dashed lines) and their interactions with various sections of ArcZ, CyaR or GcvB (blue and purple bars) or with other sRNAs and mRNAs (black bars). The CyaR sequence indicated with a blue bar is predicted to interact with two regions in GcvB (see blue bars in GcvB alignment), including the second seed sequence. A second interaction (pink bars) involves the seed sequence regions of CyaR and GcvB.

**Figure 4–figure supplement 3.**
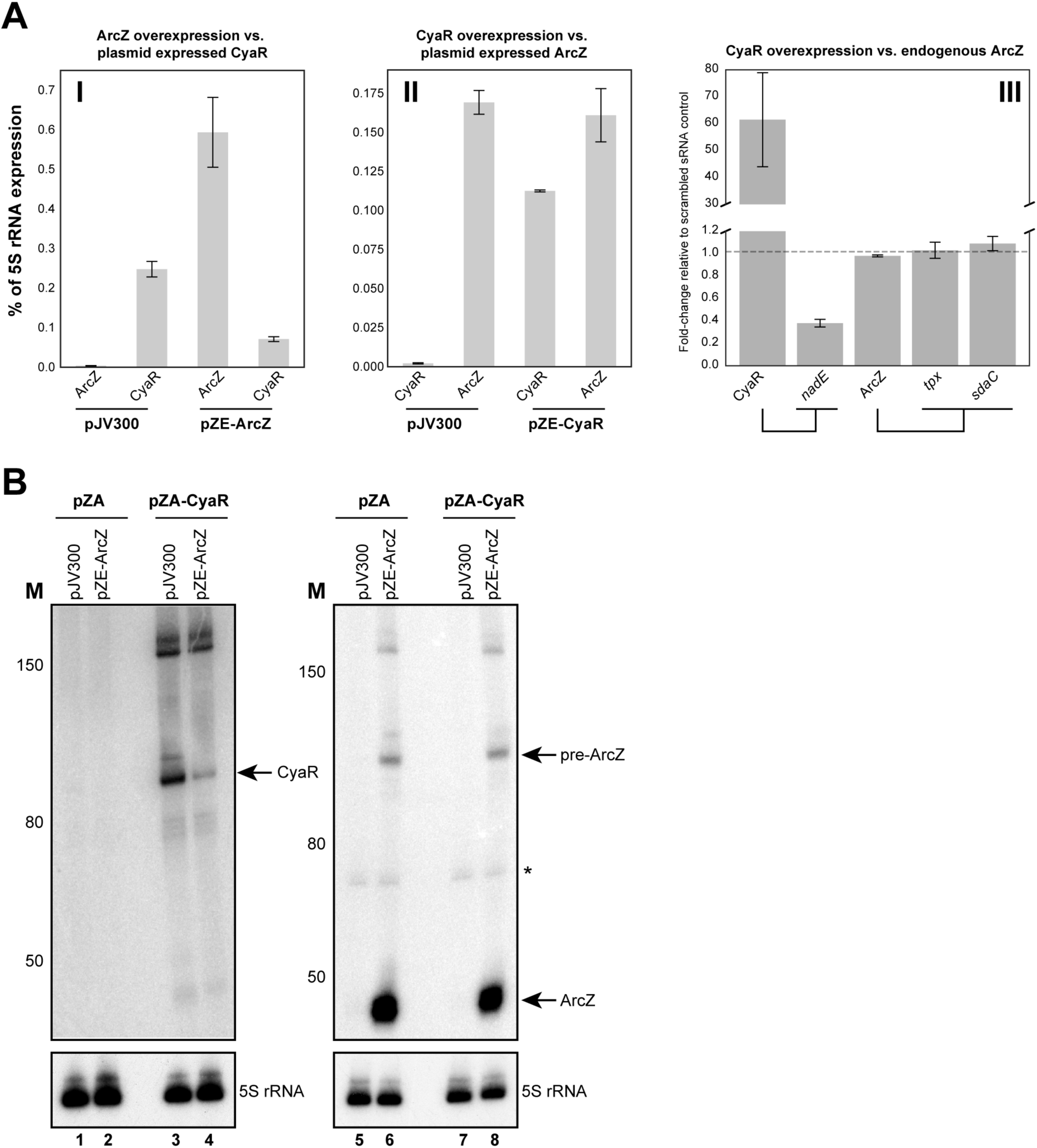
ArcZ downregulates CyaR expression. ArcZ and CyaR were overexpressed from a plasmid-borne IPTG inducible promoter (pZE-ArcZ and pZE-CyaR) and the data were compared to data from cells carrying a scrambled RNA plasmid (pJV300). The co-expressed candidate target sRNAs (expressed from pZA-derived backbone) were induced with anhydrotetracycline hydrochloride (panels I and II). The bars indicate the mean fold-change in expression relative to the level of 5S rRNA (*rrfD*) in cells with the indicated vector. In panel III endogenous ArcZ levels were measured upon over-expression of CyaR. Error bars indicate the standard error of the mean from three biological replicates and three technical replicates per experiment. A Northern blot analysis of ArcZ and CyaR. The cells containing both the empty pZA and pJV300 plasmids (lanes 1, 5, 9) do not express ArcZ and CyaR at detectable levels.

**Figure 5–figure supplement 1.**
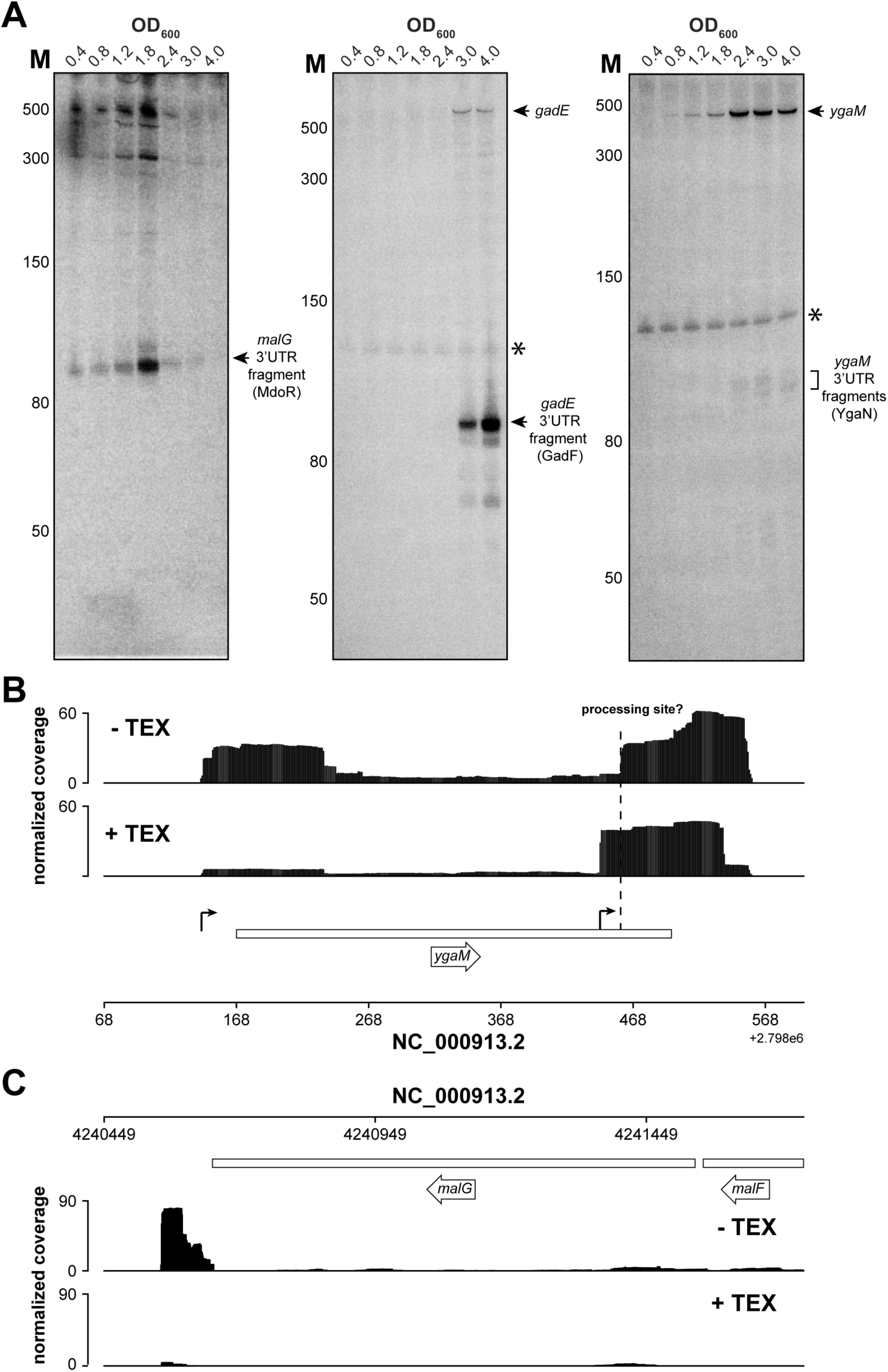
*YgaM*, *gadE* and *malG* contain sRNAs in their 3’UTRs. (**A**) Validation of *malG* 3’UTR (MdoR), *ygaM* 3’UTR (YgaN) and *gadE* 3’UTR (GadF) sRNAs by Northern blot. Total RNA extracted from cells at the indicated optical densities (OD_600_) was resolved on 8% PAA-UREA gels and subjected to Northern blotting using oligos that hybridize with the 3’UTR of the respective transcripts. The asterisk indicates cross-reactivity of the probe with the 5S rRNA. The locations of the 3’UTR-derived fragments are indicated. MdoR and YgaN are ∼110nt, whereas GadF fragment is ∼ 90nt. (**B, C**) Analysis of (Terminator 5’-Phosphate Dependent Exonuclease (TEX) RNA-seq datasets(Thomason et al., 2015) indicates that YgaN has an independent promoter, while MdoR is a degradation product of the *malEFG* operon. Genome browser tracks showing the location and normalised reads of *ygaM* and *malG* fragments in the absence of TEX (–TEX) and in the presence of TEX (+TEX). The *ygaM* and putative YgaN promoters are indicated. Independently transcribed YgaN could be further processed by RNases, at the site marked with a dashed vertical line.

**Figure 6–figure supplement 1.**
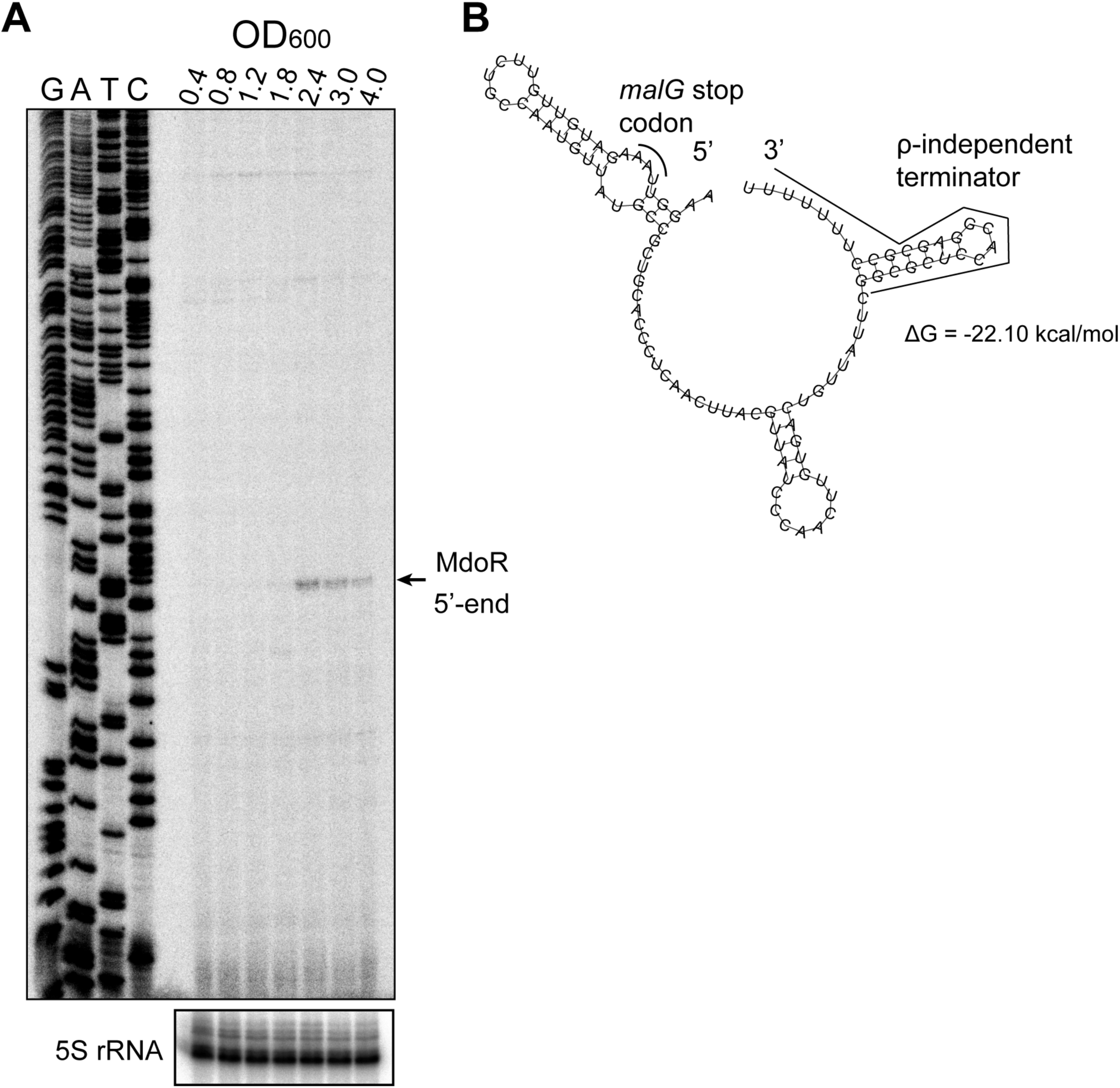
Conservation and target prediction analyses of MdoR. (A) MdoR contains conserved and variable regions. Sequence conservation analysis of MdoR in several Gram-negative bacteria species (Mafft algorithm with defaults); the arrow indicates the 5’-end of MdoR; the *malG* stop codon and Rho-independent terminator sites are highlighted; the horizontal black lines indicate the base-pairing regions of MdoR with *ptsH* (top), OhsC, *ompA* and *ompC* (bottom) in *E. coli* as predicted by CLASH combined with *in silico* folding (RNACofold). (B) MdoR is predicted to interact with its targets using two seed regions. Interaction regions within MdoR and top target mRNAs predicted by CopraRNA(Wright et al., 2014, 2013). Density plots showing the relative frequency of a specific MdoR (Left) or mRNA (Right) nucleotide position in all predicted sRNA-mRNA interactions with a p-value < 0.01 in all considered homologs. The vertical lines indicate local maxima; the aligned regions of the homologs are shown in grey, whereas the interacting regions are shown in arbitrary colors; only the top 20 representative clusters members are shown in the aligned regions, with the gene names indicated on the right.

**Figure 8–figure supplement 1.**
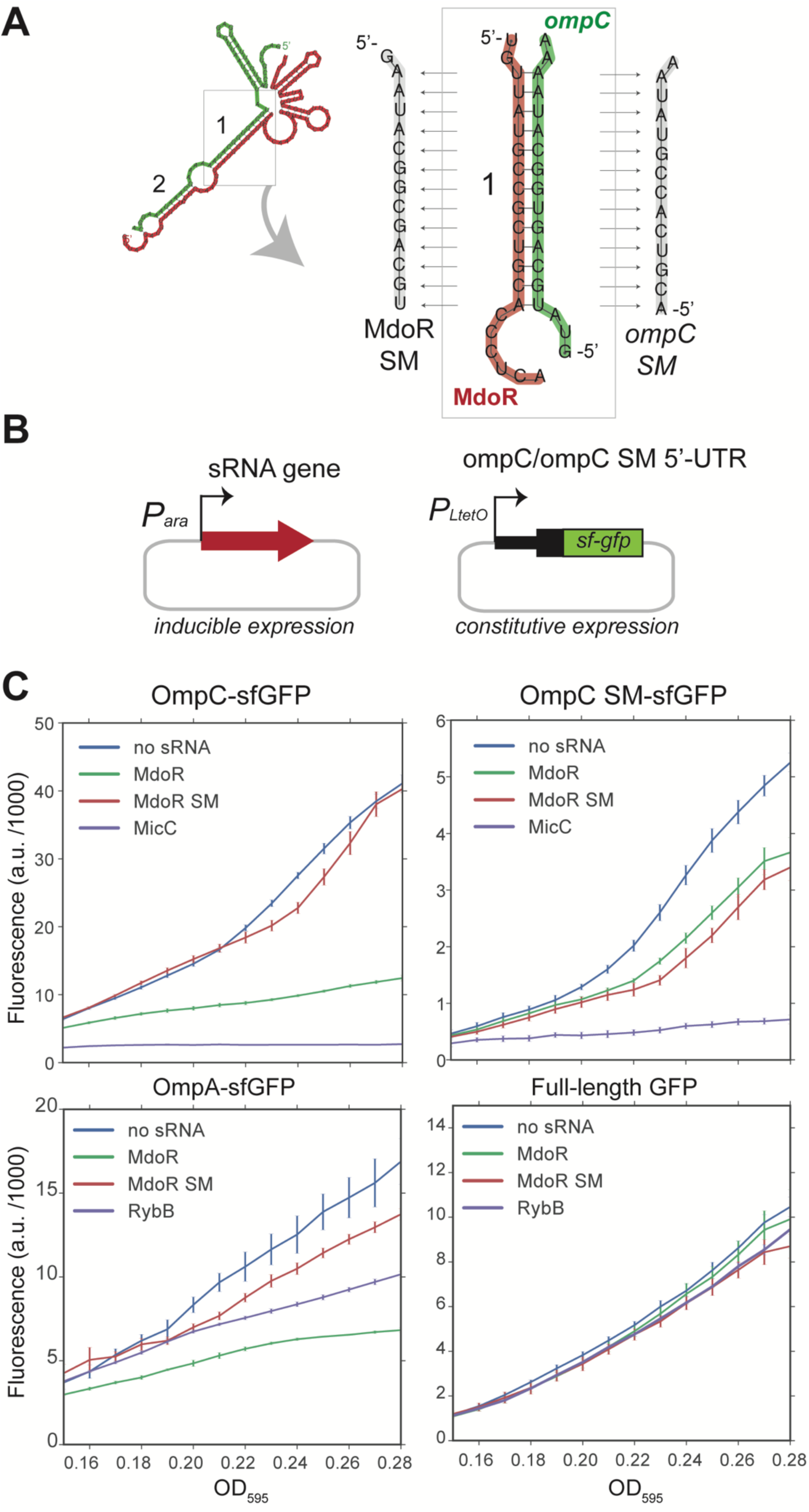
Validation of MdoR-*ompC* interaction using GFP reporters. **(A)** Design of the wild-type and mutant *ompC* constructs. The panel indicates the base-pairing region within the MdoR-*ompC* duplex that was mutated. We created an MdoR seed mutant (SM) and an *ompC* mutant in which base pairing with MdoR SM was restored. **(B)** Plasmid system used for the reporter assay: in *E. coli* TOP10 cells, low-copy plasmids constitutively overexpress target 5’UTRs fused to sfGFP and medium-copy plasmids overexpress the full-length sRNAs upon induction with L-arabinose. **(C)** MdoR downregulates expression of OmpC and OmpA sfGFP fusions. *In vivo* fluorescence measurements of OmpC, OmpC SM and OmpA sfGFP fusion proteins was measured using a Tecan plate reader system. As a negative control we included sfGFP alone in the presence or absence of sRNAs. The ‘no sRNA’ expressing strains contain the empty pBAD plasmid. The y-axis indicates fluorescence units (F.U.) reported by the plate reader. Experiments were performed in technical and biological triplicates; the fluorescence means and SEM of three biological replicates are reported. Source data for are provided as a Source Data file.

**Figure 8–figure supplement 2.**
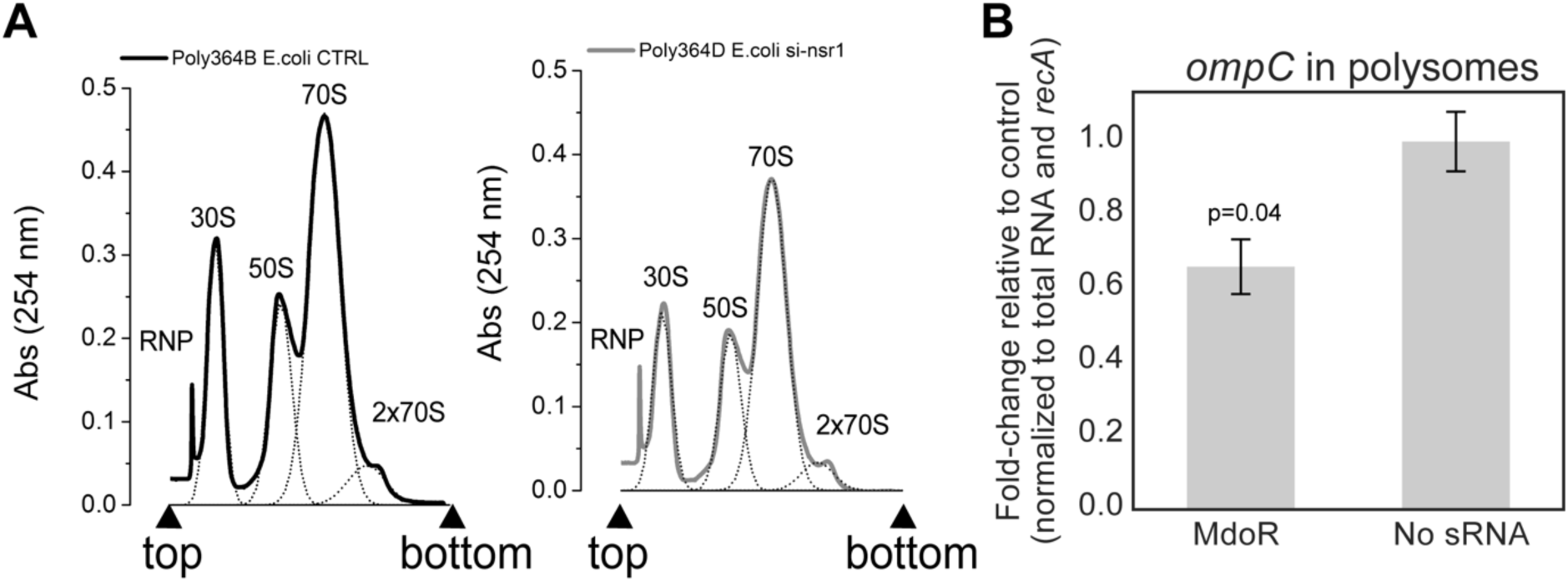
MdoR regulates *ompC* mRNA translation in *E. coli*. **(A)** Cultures at OD_600_ 0.4 overexpressing MdoR or no sRNA for 15 minutes were subjected to polysome profiling. Profiles of the polysomal (2×70S) and subpolysomal fractions obtained for the empty plasmid control and MdoR overexpression samples. **(B)** RT-qPCR analysis of the polysomal fractions: A ‘total’ fraction was obtained by mixing equal amounts/volumes of the polysomal and subpolysomal fractions and is representative of the cytosol/cell lysate content. Total RNA was extracted from all fractions (polysomal, subpolysomal and total). Expression of *ompC* in the polysomal fractions was quantified relative to the amount in ‘total’ fraction, normalized to *recA*, and calculated as fold-change relative to the control sample (y-axis). The experiments were performed in technical triplicates; the standard error of the mean (SEM) of three biological replicates fold changes are reported as error bars. Significance of the difference in *ompC* mRNA level in polysomes was assessed with a two-tailed Student’s t-test. Source data for are provided as a Source Data file.

